# Capabilities and Limitations of Tissue Size Control Through Passive Mechanical Forces

**DOI:** 10.1101/023184

**Authors:** Jochen Kursawe, Pavel A. Brodskiy, Jeremiah J. Zartman, Ruth E. Baker, Alexander G. Fletcher

## Abstract

Embryogenesis is an extraordinarily robust process, exhibiting the ability to control tissue size and repair patterning defects in the face of environmental and genetic perturbations. The size and shape of a developing tissue is a function of the number and size of its constituent cells as well as their geometric packing. How these cellular properties are coordinated at the tissue level to ensure developmental robustness remains a mystery; understanding this process requires studying multiple concurrent processes that make up morphogenesis, including the spatial patterning of cell fates and apoptosis, as well as cell intercalations. In this work, we develop a computational model that aims to understand aspects of the robust pattern repair mechanisms of the *Drosophila* embryonic epidermal tissues. Size control in this system has previously been shown to rely on the regulation of apoptosis rather than proliferation; however, to date little work has been done to understand the role of cellular mechanics in this process. We employ a vertex model of an embryonic segment to test hypotheses about the emergence of this size control. Comparing the model to previously published data across wild type and genetic perturbations, we show that passive mechanical forces suffice to explain the observed size control in the posterior (P) compartment of a segment. However, observed asymmetries in cell death frequencies across the segment are demonstrated to require patterning of cellular properties in the model. Finally, we show that distinct forms of mechanical regulation in the model may be distinguished by differences in cell shapes in the P compartment, as quantified through experimentally accessible summary statistics, as well as by the tissue recoil after laser ablation experiments.

**Author Summary:** Developing embryos are able to grow organs of the correct size even in the face of significant external perturbations. Such robust size control is achieved via tissue-level coordination of cell growth, proliferation, death and rearrangement, through mechanisms that are not well understood. Here, we employ computational modelling to test hypotheses of size control in the developing fruit fly. Segments in the surface tissues of the fruit fly embryo have been shown to achieve the same size even if the number of cells in each segment is perturbed genetically. We show that simple mechanical interactions between the cells of this tissue can recapitulate previously gathered data on tissue sizes and cell numbers. However, this simple model does not capture the experimentally observed spatial variation in cell death rates in this tissue, which may be explained through several adaptations to the model. These distinct adaptations may be distinguished through summary statistics of the tissue behaviour, such as statistics of cell shapes or tissue recoil after cutting. This work demonstrates how computational modelling can help investigate the complex mechanical interactions underlying tissue size and shape, which are important for understanding the underlying causes of birth defects and diseases driven by uncontrolled growth.

## Introduction

The mechanisms underlying tissue size control during embryonic development are extremely robust. There are many cases where the rates of proliferation, growth, or death are perturbed significantly yet patterns are maintained or repaired during later stages of development. For example, even after 80% of the material in a mouse embryo is removed, accelerated growth can give rise to correctly proportioned, albeit non-viable offspring [1]. In fruit fly embryos, overexpressing the maternal effect gene *bicoid* leads to stark overgrowth in the head region, but the excess tissue is removed during later stages of development through apoptosis (programmed cell death), leading to viable adults [2]. Tetraploid salamanders of the species *Amblystoma mexicanum* have half the number of cells as their diploid counterparts, yet are the same size [3].

The robustness of tissue size control relies on tight coordination of cellular processes including growth, proliferation, apoptosis and movement at a tissue level. However, the fundamental mechanisms underlying such coordination remain largely unknown. In particular, the mechanical implementation of tissue size control is not well understood. The regulation of cellular mechanical properties is known to play a key role during morphogenetic events, such as tissue folding, elongation and cell sorting [4, 5]. For example, upregulation of myosin II generates tension that helps to straighten compartment boundaries in the *Drosophila* wing imaginal disc [6], while controlled cell death provides the tension required for invagination during *Drosophila* leg development [7]. It has been illustrated theoretically how mechanical feedback might facilitate uniform growth in epithelia in the face of morphogen gradients [8]. Could mechanical forces also play a significant role in robustly maintaining tissue size?

To explore questions of pattern repair, we develop a computational model of a patterned epithelium, with application to the segments of the *Drosophila* embryonic epidermis (Fig. 1C). These tissues define the body plan along the head-tail axis. They are first defined during stage 6 of embryonic development and are visible as stripes in the epidermis of the larva [9]. The segments are subdivided into anterior (A) and posterior (P) compartments, which are marked by distinct gene expression patterns. In particular, cells in the P compartment express the gene *engrailed* [10] (Fig. 1D). While the initial specification and establishment of segments is relatively well studied [11], maintenance of segment identities have received much less attention. However, it is known that compartment dimensions can be robustly restored in the presence of genetic manipulations made during earlier developmental patterning events [2, 12–14]. Both the conserved epidermal growth factor receptor (EGFR) and Wnt/Wingless (WG) pathways have been implicated in regulating apoptosis to achieve pattern repair for perturbations made in each of the compartments and are known to antagonize each other [2, 14].

**Figure 1.**
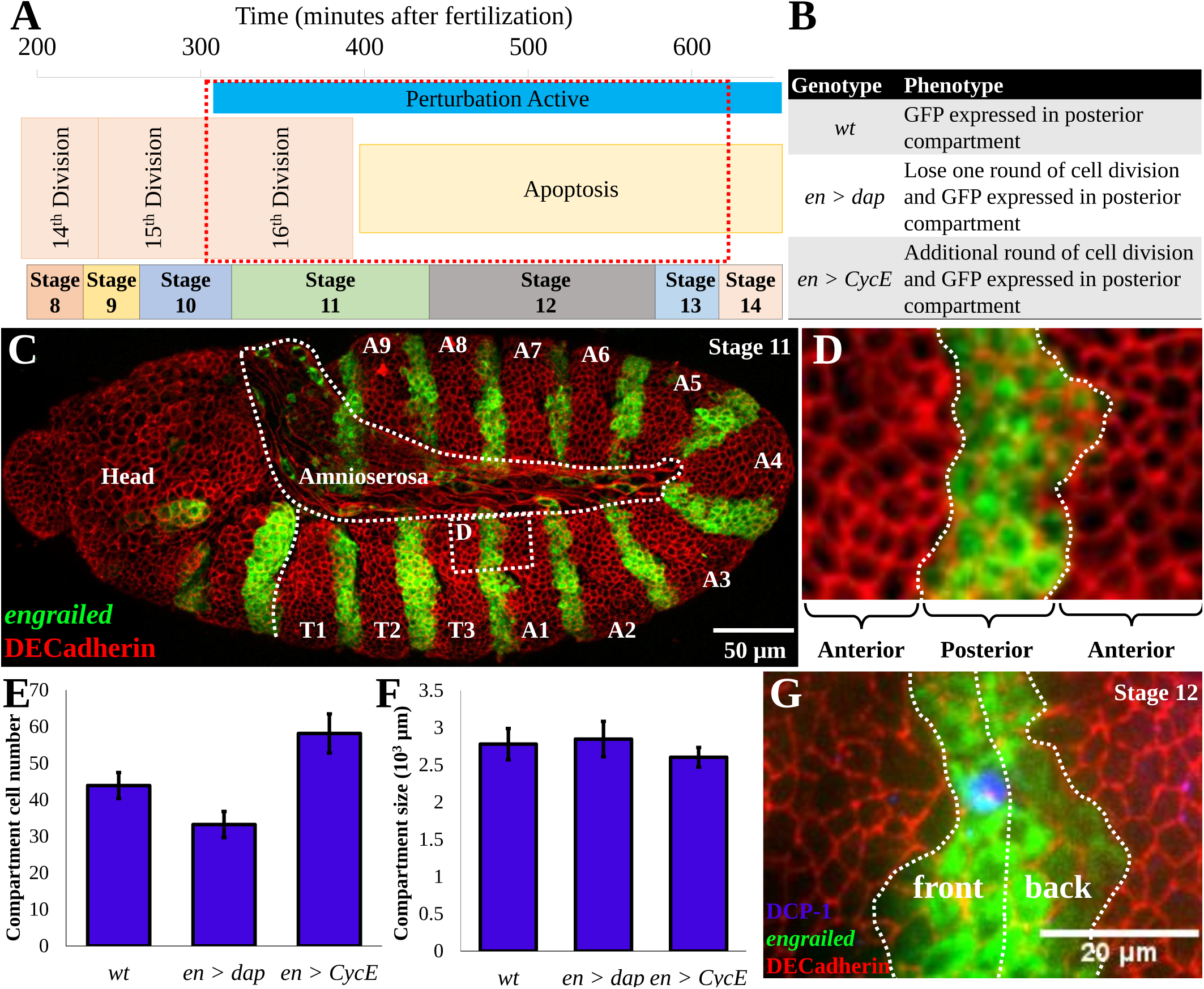
The *Drosophila* embryo as a model system for size homeostasis. (A) Specification of embryonic stages over time; the red boxed region represents the time period of simulations [19]. (B) Summary of genetic perturbations simulated in this study. The wt genotype is *engrailed>GAL4, UAS>GFP.* The perturbations are crosses between the *wt* and *UAS>CyclinE* and *UAS>dacapo* lines, respectively. (C) Stage 11 embryo expressing GFP in the posterior compartment, stained for DE-cadherin to show cell boundaries. (D) High magnification image of simulation domain. (E, F) Data extracted from [14] demonstrating that compartment dimensions are robust to manipulations that change the number of cells. (G) Cell death, indicated by cleaved *Drosophila* caspase-1 antibody staining [20], is statistically more likely to occur in the front half of the posterior compartment in *en>CycE* embryos [14].

A major strength of *Drosophila* as a model organism is the availability of genetic tools that enable the ectopic expression of gene products or RNA interference (RNAi) constructs to manipulate cell growth, proliferation and signaling in a spatio-temporally controlled manner [15–17]. For example, the bipartite GAL4-UAS system can be used to drive expression of ectopic genes in embryos through a cross of one line containing a tissue-specific enhancer driving expression of the heterologous yeast transcription factor GAL4 with a second line that activates expression of a transgene upon binding of GAL4 to the UAS promoter region. Using this approach, Parker [14] investigated P compartment size using the GAL4 driver line as the control genotype *engrailed-GAL4, UAS-GFP,* in the following referred to as *wt* (wild type). This was compared to various perturbations (Fig. 1B). In particular, these included crosses between the driver line and *UAS-CyclinE* (which we shall term *en>CycE*) and *UAS-dacapo* lines (further specified as *en>dap),* which perturbed the amount of final proliferation events towards the end of the normal range of proliferation in the epidermis (Fig. 1A).

Parker [14] observed an increase in final cell number of more than 30% (Fig. 1E, right bar) in the P compartments of *en>CycE* embryos, which exhibited an additional round of cell division. However, the size of the P compartment was much less affected by this perturbation (Fig. 1F, right bar), as measured in first instar larvae [14]. Conversely, in *en>dap* embryos that exhibited a loss of one round of cell division, Parker [14] observed a reduction in cell number of 25% while, again, the compartment size was relatively unchanged (Fig. 1E and 1F, middle bars).

Parker’s findings also suggest that epidermal growth factor receptor (EGFR) signaling, through the activating ligand Spitz, patterns apoptosis inside the P compartment. Spitz is released by a column of cells inside the anterior (A) compartment that is directly adjacent to the P compartment. Identifying cell death events through TUNEL staining [18], Parker [14] observed apoptosis much more frequently in the ‘front’ (more anterior) half of the P compartment, away from the Spitz source (Fig. 1G), than the ‘back’ half. These numbers differed by a factor of nearly 40 in *wt* [14]. Counter-intuitively, inhibiting apoptosis by expressing the caspase inhibitor p35 inside the P compartment of *en>CycE* embryonic segments resulted in compartment shrinkage by nearly 10%.

The above findings shed light on tissue size control in the *Drosophila* embryonic epidermal tissues, suggesting a reliance on the regulation of apoptosis rather than proliferation. However, the cell-level interactions governing size control remain poorly understood. In particular, potential roles of cellular mechanics in augmenting or repairing growth defects in patterned tissues remain unexplored. To address this, we develop a vertex model of an embryonic segment to test hypotheses about the emergence of size control. Comparing the model to previously published data across *wt* and genetic perturbations, we investigate the extent to which passive mechanical forces might suffice to explain the observed size control and asymmetries in cell death frequencies across the P compartment. Our results suggest that the basis of size control can rely to a significant degree on the passive mechanical responses of cells. However, the observed spatial asymmetry in cell death frequencies requires patterning of mechanical properties by inter-cellular communication. These results also provide a basis for differentiating experimentally how extracellular signaling pathways like EGFR and WG might impact cellular decision making processes through predictions of observable cellular morphologies, and tissue behaviour after cell bond ablation.

## Materials and Methods

We use a vertex model to simulate cell movement, intercalation, shape changes and apoptosis during the sixteenth round of divisions in a segment of the *Drosophila* embryonic epidermis. Vertex models were first introduced to study the structure of foams [21], and have since been applied to study a variety of epithelial tissues [6, 22–25]. For more information on vertex models and their application to epithelial morphogenesis, we refer the reader to two recent reviews [26,27].

### Equations of motion

Vertex models approximate cells in epithelial sheets as polygons. The polygons represent the cells’ apical surfaces, where most inter-cellular forces originate [4]. The terms in the model account for the mechanical effect of the force-generating molecules that accumulate in the apical surface of the cells, such as actin, myosin, and E-cadherin. Vertices correspond to adherens junctions, and their positions are propagated over time using an overdamped force equation, reflecting that adherens junctions are not associated with a momentum.

The force equation takes the form

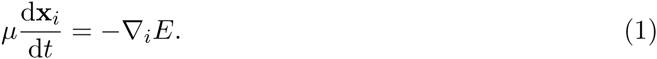

Here, *μ* is the friction strength (which we assume to take the same constant value for all vertices), *t* is time, **x***_i_* is the position vector of vertex *i*, and *E* denotes the energy of the whole system. The total number of vertices in the system may change over time due to cell division and apoptosis. The symbol ∇*_i_* denotes the gradient operator with respect to the coordinates of vertex i. The forces act to minimise a phenomenological energy function, based on the contributions thought to dominate epithelial mechanics [22]:

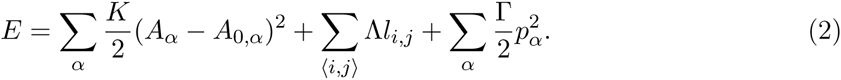

Here, the first sum runs over every cell in the sheet, *A_α_* denotes the apical surface area of cell *α* and *A*_0_*,_α_* is its preferred area, or target area. This energy term penalises deviations from a target area for individual cells, thus imposing cellular bulk elasticity. The second sum runs over all edges 〈*i, j*〉 in the sheet and penalizes long edges (we choose Λ > 0), thus representing the combined effect of E-cadherin, myosin, and actin at the binding interface between two cells. The third sum also runs over all cells, and *p_α_* denotes the perimeter of cell *α*. This term models the effect of a contractile acto-myosin cable along the perimeter of each cell [22]. The parameters *K*, Λ, and Γ together govern the strength of the individual energy contributions. Although this description of cell mechanics is phenomenological, a variety of previous studies have demonstrated its ability to match observed junctional movements and cell shapes in epithelial sheets through validation against experimental measurements [6, 22, 25].

In contrast to many previous vertex model applications, we allow the mechanical parameters Λ, Γ, and *A*_0_ to vary between cells as a consequence of underlying tissue patterning. In particular, we consider *A*_0_ to be a function of cell generation and introduce the parameter 

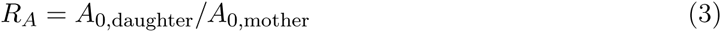
as the ratio of target areas of daughter cells to mother cells. To ensure that the target areas of all cells add up to the total size of the spatial domain, which is assumed to be fixed, we choose *R*_*A*_ = 0.5 unless stated otherwise. Throughout the study, variation of the parameter *R*_*A*_ is used to account for cellular growth of daughter cells as well as changes in total target upon division. In each simulation, the initial area of each cell, *A^s^,* equals its initial target area, 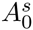, with 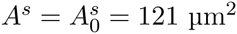 (see discussion below for the choice of length scales in the model). In Supplementary Text S1 and Supplementary Fig. S4 we analyse the extent to which deviations of cell target areas may affect the simulation results by increasing 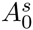. The simulated P compartment sizes and cell numbers are not strongly affected by such changes in initial condition, except for an increase in apoptosis for the *en>CycE* perturbation.

In contrast to several previous applications [22, 25] of the vertex model the spatial domain in this study is constrained due to the fact that there is no net organism growth during embryogenesis.

### Cell intercalation and apoptosis

In addition to evolving vertex positions in accordance with Eq. (1), we must allow for cell intercalation and cell removal through topological rearrangements. One such topological rearrangement is a T1 swap, which simulates cell neighbour exchanges. In a T1 swap an edge shared by two cells is removed and the cells are disconnected, while a new perpendicular edge is created that then connects the cells that were previously separated by the edge (see Fig. 2B). In our implementation T1 swaps are executed whenever the length of a given edge decreases below a threshold *l*_min_ = 0.11 μm, which is 100 times smaller than the approximate length of a cell at the beginning of the simulation. The length of the new edge, *l*_new_ = 1.05*l*_min_, is chosen to be slightly longer than this threshold in order to avoid an immediate reversion of the swap. A summary of the frequency of T1 swaps occurring in model simulations is provided in Supplementary Table S1. There are very few cell intercalation events in our simulations, with no T1 swaps observed for *wt,* in line with experimental observations of germ-band retraction [28].

**Figure 2.**
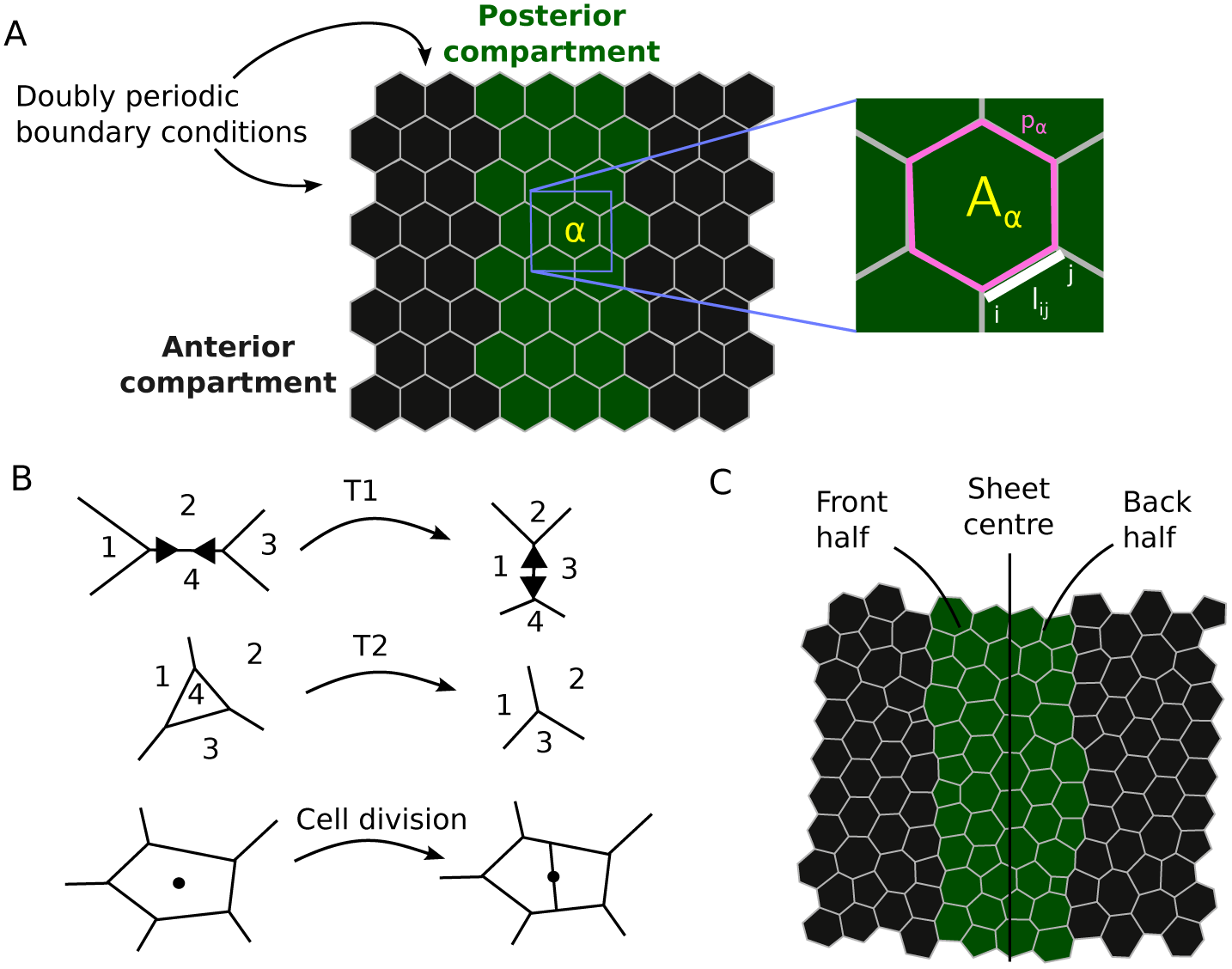
Vertex model of posterior compartment dynamics during the last division cycle in the *Drosophila* embryonic epidermis. (A) Snapshot of the initial tissue configuration for each simulation, with mechanical parameters in Eq. (2) annotated. (B) Schematic diagram of a junctional rearrangement (T1 swap), a cell removal (T2 transition), and cell division in the vertex model. Numbers indicate cell indices. (C) Snapshot of a *wt* simulation at the final time point, once all cell divisions have occurred, with annotation for the front and back halves of the P compartment. Parameter values are listed in Table 1.

A second topological rearrangement in vertex models is a T2 transition, during which a small triangular cell is removed from the tissue and replaced by a new vertex (see Fig. 2B). In our implementation any triangular cell is removed if its area drops below a threshold *A*_min_ = 0.121 μm^2^, which is 100 times smaller than the initial area of each cell. The energy function (2) in conjunction with T2 transitions can be understood as a model for cell removal: cells are extruded from the sheet by a T2 transition if they become mechanically unstable. Note that we do not discriminate between cell removal by cell death or by delamination, since this distinction is immaterial for our purposes. However, delamination has been shown to provide an alternative way of cell removal from epithelial sheets that is distinct from apoptosis [29]. Rates of cell removal predicted by previous vertex model applications have coincided with experimental measurements in the *Drosophila* wing imaginal disc [22] and notum [29].

### Cell proliferation

All simulations start with 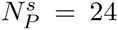 cells in the P compartment and 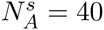 cells in the anterior compartment, to approximately match observed cell numbers [14] and to ensure that there are similar amounts of anterior tissue on either side of the P compartment.

In the case of a *wt* embryonic segment each cell divides once, with cell cycle times drawn independently from the uniform distribution on 0 to 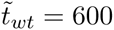 time units. This corresponds to the duration of the sixteenth division cycle in the epidermis, which occurs during late stage 10 and early stage 11 and takes roughly 50 minutes [19]. After the round of divisions is complete, the system is allowed to relax for 200 more time units, corresponding to a total simulation time of *t*_*wt*_ = 800 time units.

For an *en>CycE* embryonic segment, the first round of divisions is implemented as for *wt,* but each cell in the P compartment then has a probability *P*_*CycE*_ = 0.54 of dividing a second time once the first round of divisions is complete, with cell cycle times drawn independently from the uniform distribution from 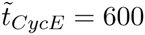 to 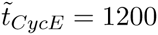 time units. This probability is inferred from published data on the *en>CycE+p53* perturbation [14]; in this case apoptosis is blocked, allowing us to infer the average number of cell division events. The second period of 600 time units corresponds to the duration of the ectopic divisions in the *en>CycE* embryos, which occur during late stage 11 and early stage 12 [14]. After the second round of divisions is complete, the system is allowed to relax for 200 more time units, corresponding to a total simulation time of *t*_*CycE*_ = 1400 time units.

For an *en>dap* embryonic segment, each cell in the P compartment has a fixed probability *P*_*dap*_ = 0.6 of not participating in the single round of divisions. This probability is inferred from published data on the *en>dap* perturbation [14]. As with *wt,* divisions occur during the first 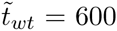 time units, after which the system is allowed to relax for 200 more time units, corresponding to a total simulation time of *t*_*wt*_ = 800 time units.

These simulation times are chosen such that the system is at quasi-steady state at each time point. This quasi-steady state assumption is commonly used in vertex models [6, 22, 29, 30] and reflects the fact that the times associated with mechanical rearrangements (seconds to minutes) are an order of magnitude shorter than typical cell cycle times (hours) [22].

At each cell division event, a new edge is created that separates the newly created daughter cells. The new edge is drawn along the short axis of the polygon that represents the mother cell [31]. The short axis has been shown to approximate the division direction (cleavage plane) of cells in a variety of tissues [32], including the *Drosophila* wing imaginal disc [33]. The short axis of a polygon crosses the centre of mass of the polygon, and it is defined as the axis around which the moment of inertia of the polygon is maximised. Each daughter cell receives half the target area of the mother cell upon division unless stated otherwise.

### Geometry and boundary conditions

In order to simulate the subsections of the P compartment we consider a spatial domain comprising two adjacent cell populations, the cells in the P compartment and parts of the adjacent tissue in the anterior compartment on either side of it. Sample simulation images are shown in Fig. 2A and Fig. 2C. For simplicity, we assume that cells initially have regular hexagonal shapes. We analyse the sensitivity of P compartment sizes and cell numbers to this choice of initial condition in Text S1 and Figure S3.

Motivated by the repeated pattern of A and P compartments along the anterior-posterior axis of the embryo, as well as by the fact that single P compartments stretch farther dorsoventrally than the simulated region, doubly periodic boundary conditions are applied (Fig. 2A). These boundary conditions keep the simulated region of interest at a fixed size. Compartment size changes are analysed as changes in the relative proportions of the anterior and posterior compartment within the simulated region.

An analysis of the sensitivity of P compartment sizes and cell numbers to this choice of boundary condition is provided in Text S1 and Figure S1. The precise choice of boundary condition imposed in the model simulations does not significantly affect predicted compartment sizes and cell numbers.

To enable comparison of cell death rates in the front and back halves of the P compartment (see Fig. 1G), a cell is defined to be in the front or back half if its centroid is located to the anterior (‘left’) or posterior (‘right’) side of the centre of the tissue, respectively. The tissue centre is defined to be the horizontal midpoint of the sheet at time *t* = 0 and is held fixed at all times.

When computing measures of cell shape in our analysis of simulation results, we define the area and perimeter of a cell to be those of the associated polygon in the vertex model, while ‘cell elongation’ is defined as the square root of the ratio of the largest to the smallest eigenvalues of the moment of inertia of that polygon. This latter measure provides a robust way to measure elongations of arbitrary shapes [31] and is comparable to the ratio of the lengths of the long and short axis of the best fit ellipse to a cell.

### Compartment boundary line tension

Unless stated otherwise, the line tension along the compartment boundaries is set to Λ*_b_* = 2Λ, twice the value of the line tension in the remainder of the tissue. High tension along compartment boundaries is known to promote cell sorting and boundary straightness [6, 30], and the presence of myosin cables that can generate this tension between A and P compartments in the *Drosophila* embryonic epidermis has been reported [34]. Figure S2 shows that while the increase in line tension along compartment boundaries does affect the straightness of the boundary between A and P compartments in the model simulations, it does not significantly affect compartment sizes or cell numbers.

### Incorporating mechanical asymmetry

To investigate the consequences of asymmetries in cell mechanical properties on P compartment size control and patterning of apoptosis, we consider three distinct cases.

In the first case, we allow for asymmetry in cell target areas in the P compartment. This is implemented by modifying the target area of each cell in the P compartment to take the form

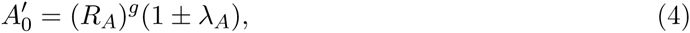
where *R*_*A*_ = 0.5 as listed in Table 1, *g* ∈ {0, 1, 2} denotes the generation of the cell, and the – and + signs apply to cells located in the front and back halves of the compartment, respectively. We refer to the parameter λ*_A_* as the *area asymmetry.*

In the second case, we allow for asymmetry in line tensions in the P compartment. This is implemented by modifying the line tension of each cell-cell interface (edge) inside the P compartment to take the form

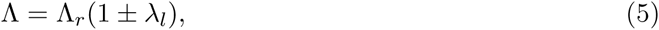
where Λ*_r_* is the value of the line tension when no asymmetry is imposed. The + sign applies to all edges between P compartment edges whose midpoint is the front half of the compartment, while the – sign applies to all edges whose midpoint is in the back half of the compartment. We refer to the parameter λ*_l_* as the *line tension asymmetry.*

In the third case, we allow for asymmetry in perimeter contractility in the P compartment. This is implemented by modifying the perimeter contractility of each cell in the P compartment to take the form 

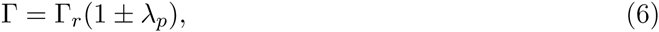
where Γ*_r_* is the value of the perimeter contractility when no asymmetry is imposed, and the + and – signs apply to cells in the front and the back halves of the P compartment, respectively. We refer to the parameter *λ_p_* as the *perimeter asymmetry.*

The asymmetry parameters are all fixed at 0 in Figs. 3, S1 and S2, and are varied in Figs. 4, 5 and 6.

**Figure 3.**
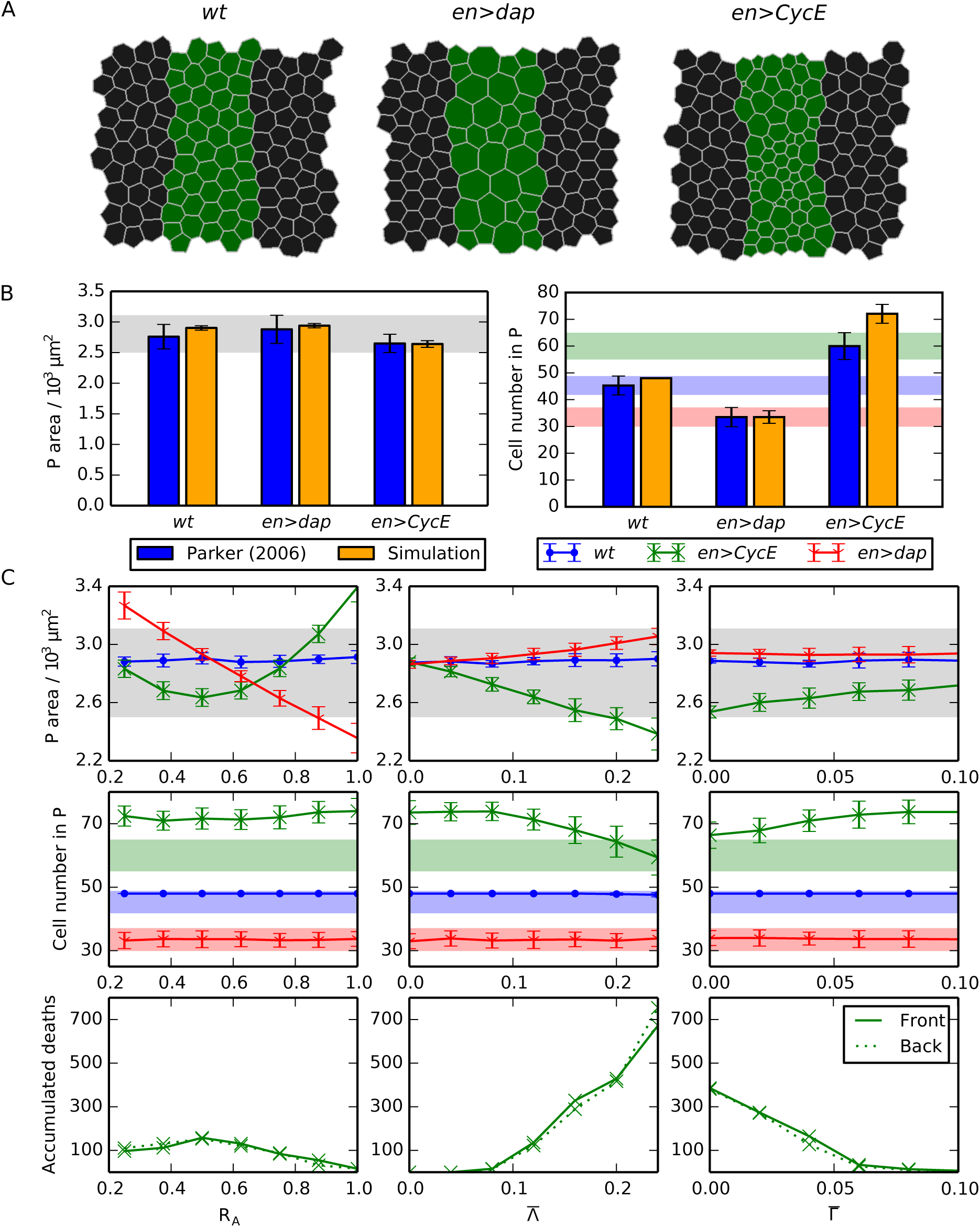
Compartment size control can emerge from passive mechanical forces. (A) Snapshots of *wt, en>dap* and *en>CycE* simulations, each following the final round of division. Parameter values are listed in Table 1. (B) Comparison of simulated P compartment areas and cell numbers with observed values [14]. Mean values from 100 simulations are shown and error bars are standard deviations. (C) Variation of P compartment area (upper row) and cell number (middle row), and of the number of accumulated cell deaths in the *en>CycE* perturbation over 100 simulations in the front and back halves of the P compartment (lower row), as each mechanical parameter is varied individually, holding all other parameters at their values listed in Table 1. Shaded areas in (B) and (C) mark the ranges of experimentally observed values and are added for reference (see main text for details).

**Figure 4.**
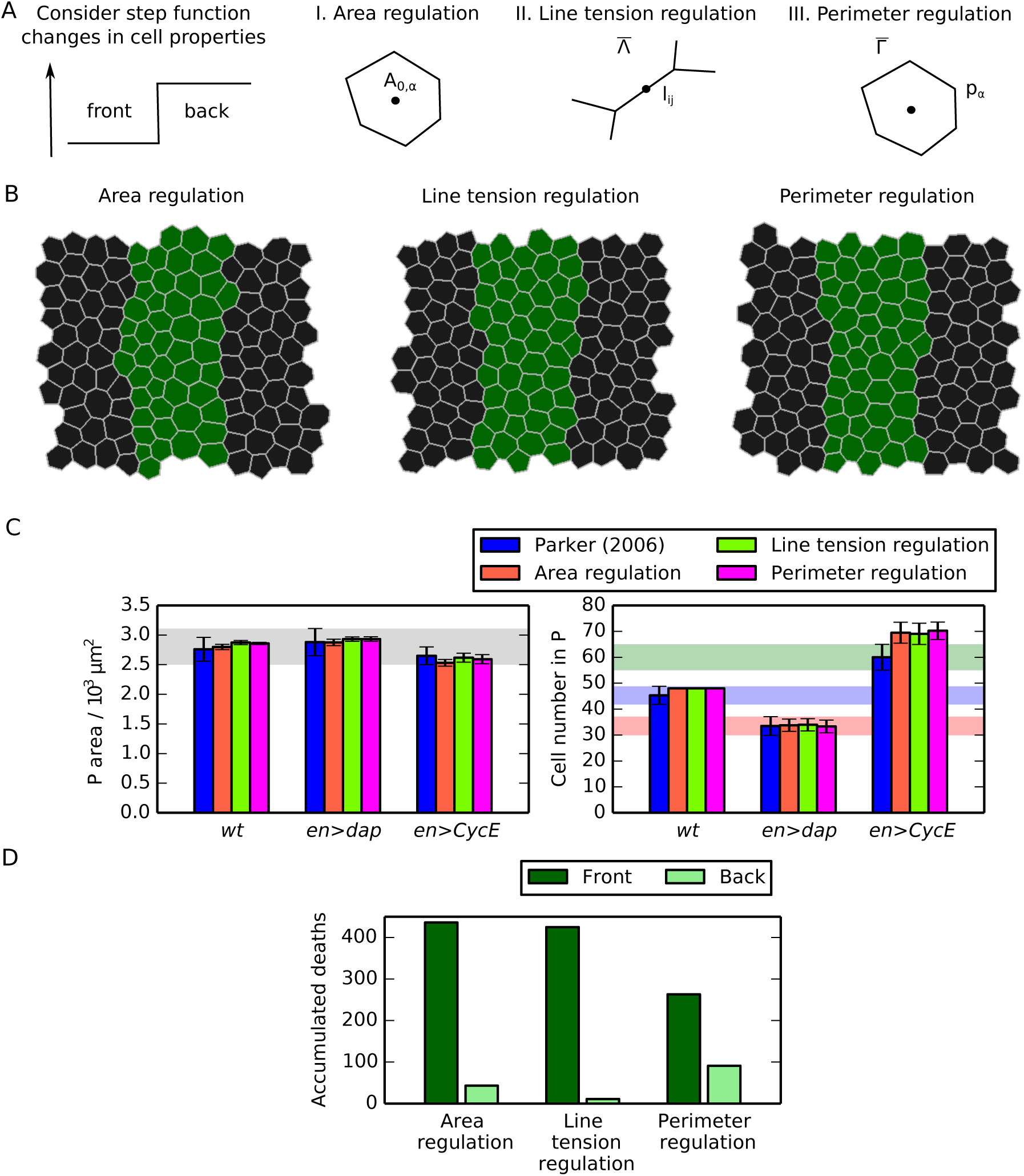
Spatial regulation of mechanical cell properties can induce asymmetry of cell death occurrence inside posterior compartments. (A) Schematic of the distinct forms of mechanical asymmetries considered in this work. (B) Snapshot of final configuration of simulations for each considered perturbation. (C) Comparison of P compartment areas and cell numbers for each of the considered perturbations with experimental values. Mean values from 100 simulations are shown and error bars are standard deviations. Parameter values are given in Table 1 and in the main text. Shaded areas mark the ranges of experimentally observed values and are added for reference and comparison with Figs. 3, S1 and S2. (D) Comparison of accumulated number cell death over 100 simulations in the front and back halves of the P compartment for each of the considered perturbations.

**Figure 5.**
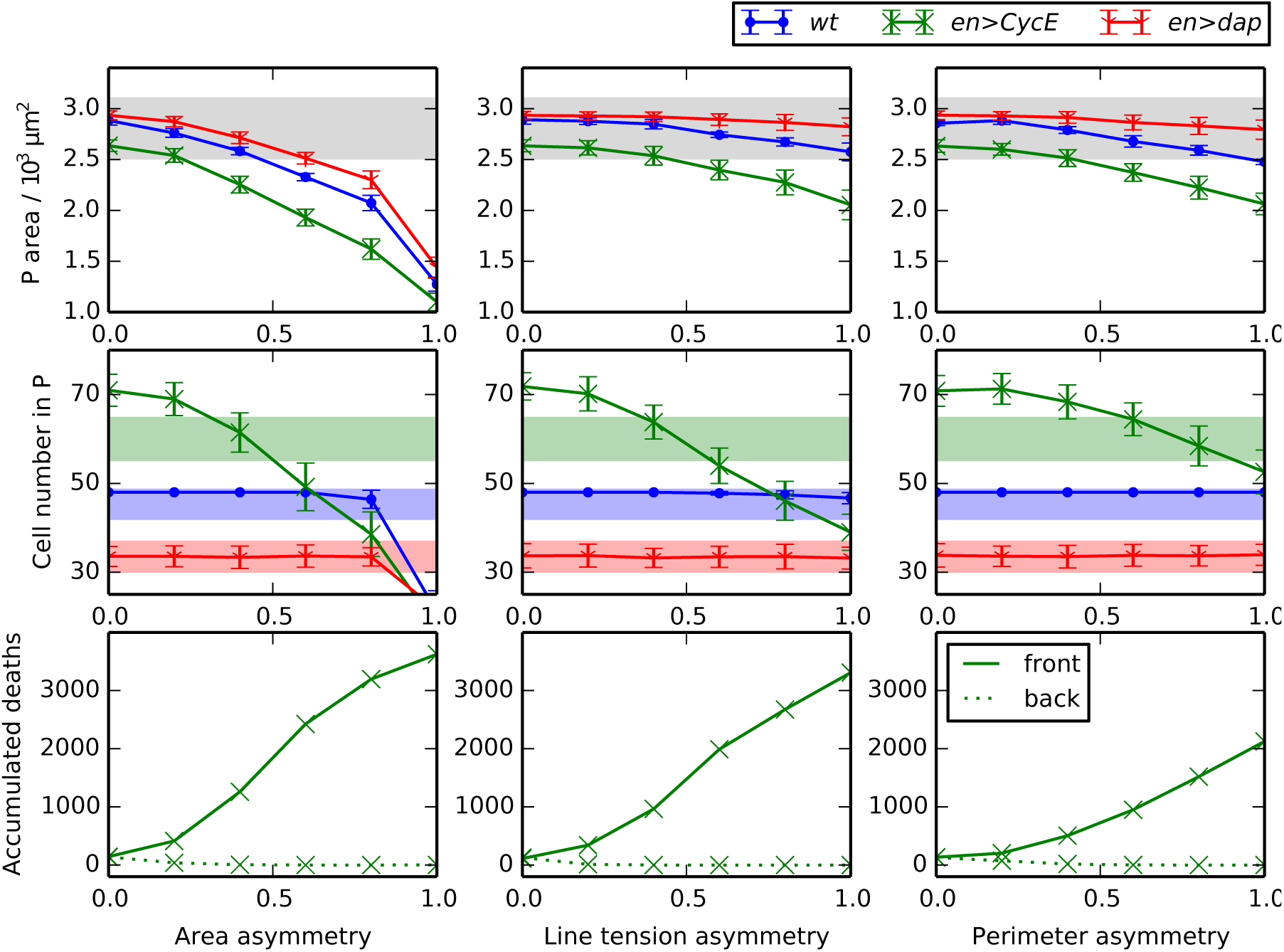
Sensitivity of P compartment size and cell number to asymmetry. Variation of P compartment area (upper row) and cell number (middle row), and of the number of accumulated cell deaths over 100 simulations in the front and back halves of the P compartment (lower row), as the asymmetry parameters *λ_A_*, *λ_l_*;, and *λ_p_* individually while holding all other parameters at their values listed in Table 1. Shaded areas are added for comparison with Figs. 3 and 5.

**Figure 6.**
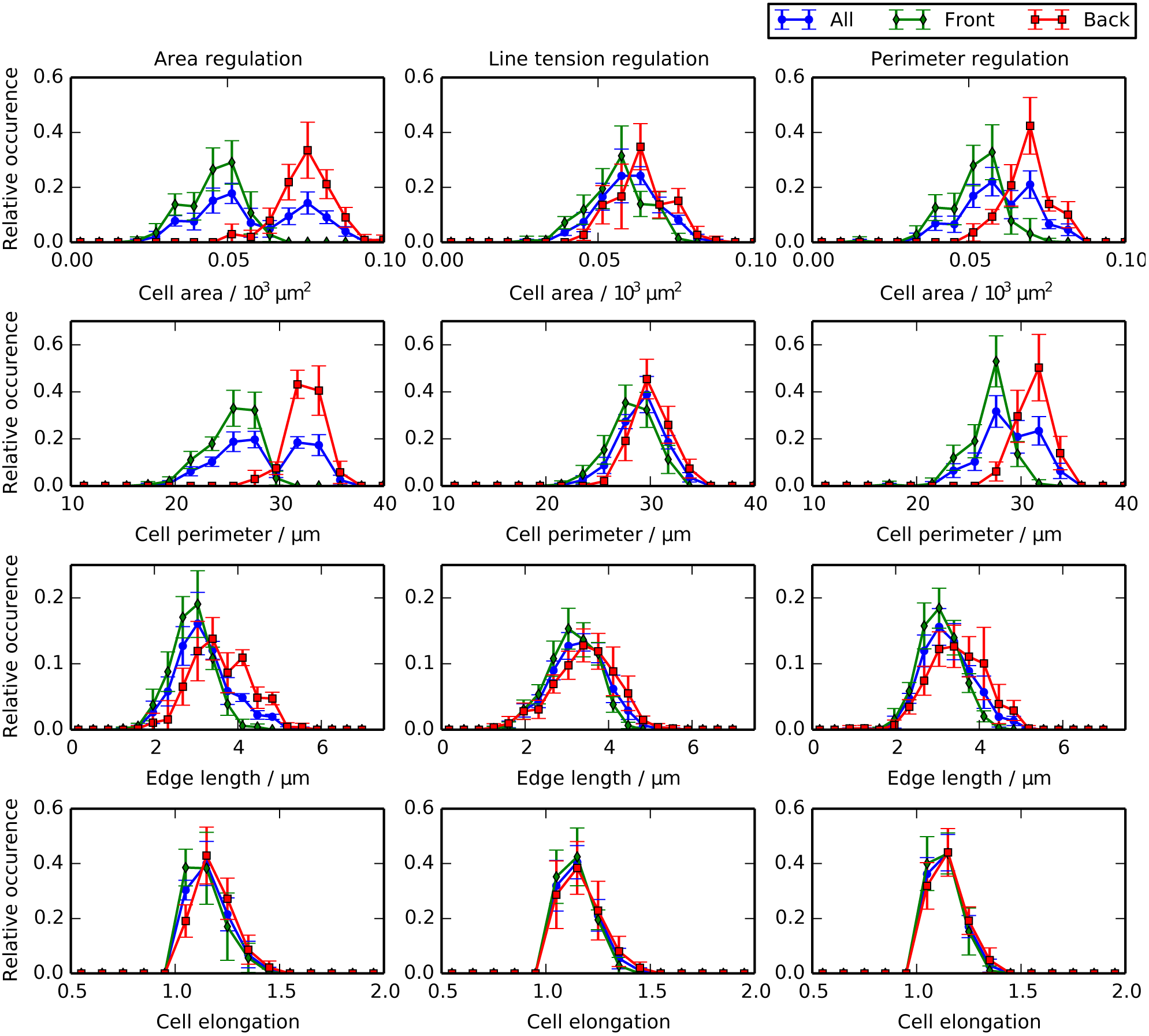
Differential growth and mechanical regulation generate distinct distributions of cell shapes. Distributions of cell areas (row 1), cell perimeters (row 2), cell edge lengths (row 3), and cell elongations (row 4) for the *wt* simulations of each scenario of cellular asymmetry. We distinguish distributions for all cells in the posterior compartment (‘All’), for cells the cells in the front half (‘Front’), and cells the back half (‘Back’).

### Numerical implementation

Prior to solving the model numerically, we non-dimensionalise it. Non-dimensionalising reduces the number of free parameters in the system and facilitates comparison of parameter values to previous implementations of the vertex model [22]. Rescaling space by the characteristic length scale *L* and time by the characteristic time scale *T*, equations (1) and (2) become 

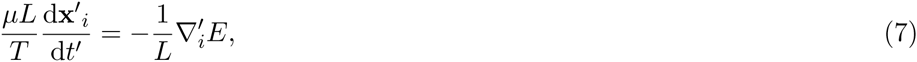

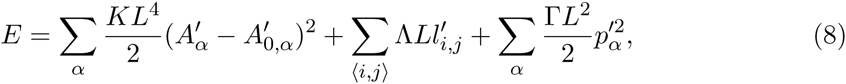
where 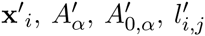 and 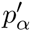 denote the rescaled *i*^th^ vertex positions, the rescaled cell area and cell target area, and the rescaled cell perimeter, respectively. The symbol 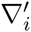 denotes the gradient with respect to the rescaled *i*^th^ vertex position. Multiplying the first equation by *T/μL,* we obtain

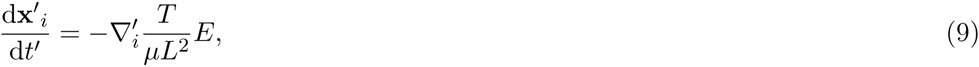

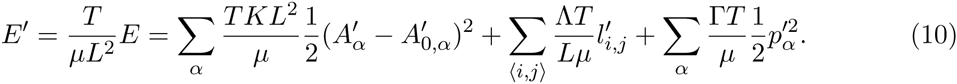

Finally, by introducing the time scale *T* = *μ*/*KL*^2^, and the rescaled mechanical parameters 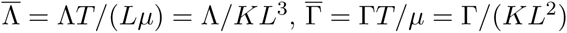 the non-dimensionalised equations read

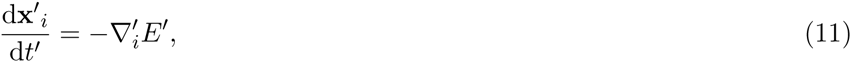

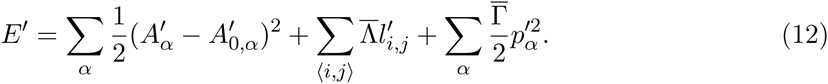

We choose the characteristic length scale *L* = 11 μm such that *L*^2^ is the mean cell area in the P compartment at the start of the simulation period, i.e. 121 μm^2^; the P compartment occupies a total area of 2.76×10^3^ μm^2^ [14] and is initialized with 24 cells. The precise value of the characteristic time scale *T* depends on tissue properties (*μ* and *K*) and could be inferred from the duration of vertex recoil after laser ablation experiments, for example. In the non-dimensionalised model, cell shapes are governed by the rescaled target area of each cell and the rescaled mechanical parameters, 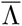 and 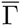. For these parameters we use previously proposed values [22], unless stated otherwise. A complete list of parameters used in this study is available in Table 1.

**Table 1.** Description of parameter values used in our simulations.

| Parameter | Description | Value | Reference |
| --- | --- | --- | --- |
| $\bar{\Lambda}$ | Adhesion parameter between cells in same compartment | 0.12 | [22] |
| $\bar{\Lambda}_r$ | Reference adhesion parameter for asymmetry simulations | 0.12 | [22] |
| $\bar{\Lambda}_b$ | Adhesion parameter between cells in different compartments | $2 \times 0.12$ | [6] |
| $\bar{\Gamma}$ | Cell perimeter contractility | 0.04 | [22] |
| $\bar{\Gamma}_r$ | Reference perimeter contractility for asymmetry simulations | 0.04 | [22] |
| $\Delta t'$ | Time step | 0.01 | [31] |
| $A'_{\min}$ | T2 transition threshold | 0.001 | [31] |
| $l'_{\min}$ | T1 swap threshold | 0.01 | [31] |
| $l'_{\text{new}}$ | Distance between new edge nodes after swaps | $1.05 l'_{\min}$ | [31] |
| $t_{wt}$ | End time of the simulation for *wt* and *en>dap* | 800 | – |
| $t_{CycE}$ | End time of the simulation for *en>CycE* | 1400 | – |
| $\hat{t}_{wt}$ | Time when first round of divisions finishes | 600 | – |
| $t_{CycE}^s$ | Time when second round of divisions starts in *en>CycE* | 600 | – |
| $\hat{t}_{CycE}$ | Time when second round of divisions finishes in *en>CycE* | 1200 | – |
| $A'^s$ | Initial cell area | 1.0 | [22] |
| $A_0^s$ | Initial cell target area | 1.0 | [22] |
| $N_P^s$ | Initial cell number inside P compartment | 24 | [14] |
| $N_A^s$ | Initial cell number inside A compartment | 64 | [14] |
| $P_{dap}$ | Probability for P cells to not divide in *en>dap* | 0.6 | [14] |
| $P_{CycE}$ | Probability for P cells to divide twice in *en>CycE* | 0.54 | [14] |
| $R_A$ | Ratio between target areas of mother cells and daughter cells | 0.5 | – |
| $\lambda_A$ | Area asymmetry | 0.0 | – |
| $\lambda_l$ | Line tension asymmetry | 0.0 | – |
| $\lambda_p$ | Perimeter asymmetry | 0.0 | – |

For parameter values for which no reference is given, please see main text for details on how these values were estimated.

To solve Eqns. (11) and (12) numerically we use an explicit forward Euler scheme:

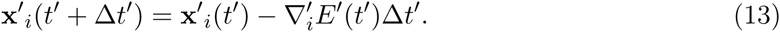

The time step used in the forward Euler scheme is 0.01 rescaled time units and is manually chosen to ensure that the numerical scheme converges and that a further reduction in the time step does not change the results.

We implement the model within Chaste, an open source C++ library that provides a systematic framework for the simulation of vertex models [31, 35]. All code used to implement model simulations and to generate results presented in this work is provided (see Software S1).

## Results

We first analyse the extent to which passive mechanical forces can lead to stable tissue size control as observed in [14]. We then investigate the effect of spatial regulation of cellular mechanical properties on P compartment sizes, cell numbers, and cell death locations.

### Compartment size control can emerge from passive mechanical forces

As an initial study, we analyse simulations where compartment size is governed solely by passive mechanical properties of individual cells, and no further regulatory mechanism for size control is assumed. In particular, all cells in the tissue are specified to have the same mechanical properties, with the exception of interfaces shared by cells at the compartment boundary. As we shall show, such passive mechanical interactions are sufficient to explain the robustness of compartment size to hyperplastic manipulations.

Fig. 3A shows snapshots of individual simulations of *wt, en>dap* and *en>CycE* embryonic segments. We observe cells that are larger but fewer in number in *en>dap* than in *wt,* while the *en>CycE* compartment contains more smaller cells. Generating statistical distributions by running 100 simulations in each case, we obtain the summary statistics visualized in Fig. 3B-C. To allow for comparison with observed values we superimpose on each panel in Fig. 3B-C either the upper and lower bounds in observed P compartment areas [14] across the three perturbations (shaded gray) or the upper and lower limits in cell numbers for each perturbation separately (blue, green, red for *wt, en>CycE,* and *en>dap,* respectively). We do not plot the distinct shaded regions in the case of P compartment areas since the regions for individual perturbations overlap. Fig. 3B shows that, for *wt* and *en>dap,* the average P compartment sizes and cell numbers at the end of the final round of divisions predicted by the model closely match observed values. The difference in cell number between simulation and experiment for *en>CycE* is statistically significant (17%), indicating that the model underestimates the number of cell deaths in this perturbation.

These simulation results were achieved using literature values of the parameters 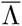 and 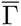 [22], and by assigning daughter cells to have half the target area of their mother cells (*R*_*A*_ = 0.5). Although the model is a drastic simplification of epithelial compartment size homeostasis, the *in silico* results provide a close match to experimental values without any parameter tuning. The model thus provides a simple explanation for the emergence of P compartment size control [14]: size control can be achieved through passive mechanical forces without any further regulation of cellular properties through signaling gradients.

To explore how robust the observed size control is to the model parameters, we performed a single parameter sensitivity analysis while fixing the remaining parameters at their values listed in Table 1 (Fig. 3C). For most parameter values considered, the simulation results fall within the bounds of experimentally observed values, except for values of the target area ratio *R*_*A*_ smaller than 0.4 and larger than 0.9, and for values in Λ larger than 0.2.

Focusing on the results of *en>CycE* simulations, the model exhibits some counter-intuitive behaviour. In particular, uniformly low perimeter contractility, 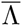, or high line tension, 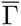, leads to mechanically induced P compartment shrinkage. In addition, an increase or decrease of *R*_*A*_ away from 0.5 will increase compartment sizes for the *en>CycE* perturbation. We may interpret these results as follows.

Mechanically induced P compartment shrinkage can be understood as a result of the balance between the energy terms in equation (2). The perimeter contractility and line tension terms act to minimise edge lengths and perimeters of cells. These force contributions can be counteracted by the area term, which acts to keep the cell close to its target area, or by stretching forces exerted by neighbouring cells. Upon division, a new edge is created, which adds an inward contractile force that any expansive forces must counteract. Therefore, daughter cells occupy a smaller area than their mother cell once they reached mechanical equilibrium. The observation that an increased rate of cell division leads to tissue shrinkage is counter-intuitive, yet not unrealistic; data from [14] for *en>CycE* and *en>CycE+p53* embryonic compartments show a similar trend, in which the more cells are present, the smaller the compartment area. Inhibition of cell death in the *en>CycE+p53* leads to more cells, but smaller compartments. Further, this counter-intuitive experimental result, which cannot be explained by a simpler hypothesis where EGFR signaling leads to size control through direct patterning of apoptosis and growth, may be explained by a simple mechanical argument.

A similar mechanism explains the dependence of the size of the *en>CycE* compartment on the target area ratio, *R_A_.* Mitosis induced shrinkage is a result of the perimeter contractility and line tension terms in the mechanical model. If we choose a value for *R*_*A*_ that is not equal to 0.5, then the target areas of all cells will no longer add up to the total area of the tissue, and more cells have areas that are far away from their actual target areas. This increases the absolute value of the area elasticity term in the energy equation, and hence reduces the relative strength of the perimeter contractility and line tension terms. As the relative strength of these two terms decreases, the extent of mitosis-induced shrinkage is also reduced. In the case *R*_*A*_<0.5, the additional line tension and perimeter force due to the new edge during division are not strong enough to stretch the cells surrounding the division further away from their target area, and if *R*_*A*_>0.5 the forces originating from the new edge are not strong enough to further oppose the strength of the target area terms of the new cells. Hence, mitosis-induced shrinkage occurs only if *R*_*A*_ ≈ 0.5. In our simulations, P compartment size is relatively robust to the value of *R_A_,* despite the fact that the areas of many cells differ widely from their target values. The bulk elasticity energy term in equation (2) varies quadratically with deviations between cell area and cell target area. Thus, one might expect significant changes in P compartment areas or cell numbers when target areas are perturbed upon proliferation. Our simulation results suggest that P compartment areas or cell numbers are not affected by such changes in total tissue energy.

A further counter-intuitive result shown in Fig. 3C is that increasing the line tension parameter 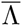 and increasing the perimeter contractility parameter 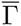 have opposing effects on P compartment size in the *en>CycE* perturbation. Increasing line tension results in a stronger contractile force on the cell, resulting in more T2 transitions and hence a smaller P compartment (Fig. 3C, central panel). In contrast, although increasing perimeter contractility also results in a stronger contractile force for each cell, in this case the mechanical interactions between adjacent cells (a contracting cell acts to stretch its neighbours) result in fewer T2 transitions and hence a larger P compartment.

All the observed changes in P compartment sizes and cell numbers remain within experimentally measured values (Fig. 3, shaded regions), the exception being the P compartment cell numbers for the *en>CycE* perturbation. The discrepancy between observed values and *in silico* results for the P compartment cell numbers in *en>CycE* is insensitive to parameter variation. The robustness of the simulation results in Fig. 3B to parameter values provides further confirmation that size control is a natural outcome of passive mechanical cellular interactions in our model. Size control is preserved in the face of small amounts of cell growth or shrinkage (variations in *R*_*A*_) or perturbations of cellular mechanical properties (variations in 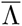 and 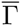).

However, this model fails to capture the observed asymmetry in cell death locations, as measured by the ratio of accumulated cell death occurrence between the front and the back half of the P compartment across multiple embryos. The third row of Fig. 3C shows that the total number of cell deaths across 100 simulations is the same between the front half and the back half of the P compartment. Here we only plot the cell death occurrences of the *en>CycE* simulations, since no cell deaths were observed in any *wt* or *en>dap* simulations. This is in close agreement with experimental results [14], where only 0.7 (*wt*) or 0.2 (*en>dap*) cell deaths where identified by TUNEL staining per embryo.

We draw two main conclusions from the simulations presented in Fig. 3: (i) mechanical interactions between identical cells can explain robust size control of all considered genetic perturbations (*wt*, *en>CycE, en>dap),* even if the parameters are varied significantly; (ii) passive mechanical interactions of cells with uniform mechanical properties cannot explain the observed asymmetry in cell death occurrence, nor completely recapitulate the changes in cell numbers for the *en>CycE* perturbation.

### Spatial patterning of cell death emerges from differential growth or differential mechanical regulation

We next use the model to analyse how asymmetries in cellular mechanical properties across the P compartment may lead to the observed spatial patterning of apoptosis. We consider three cases (Fig. 4A): (i) ‘area regulation’, which refers to patterning of the cell target areas *A*_0_,*_α_* through the parameter *λ_A_*; (ii) ‘line tension regulation’, which refers to patterning of the line tension 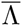 through the parameter *λ_l_*; and (iii) ‘perimeter regulation’, which refers to the patterning of the perimeter contractility 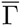 through the parameter *λ_p_*. These parameters are defined in the *Materials and Methods* section. The ‘area regulation’ scenario can be interpreted as a patterned growth scenario, whereas the ‘line tension regulation’ and ‘perimeter regulation’ scenarios correspond to patterning of cellular mechanical properties. The biochemical process leading to such patterning could, for example, be Spitz-mediated EGFR-activation; this pathway has previously been identified to affect cell properties in the P compartment by Parker [14].

Fig. 4B-D shows the impact of small amounts of asymmetry on P compartment dynamics. In each of the cases (i)-(iii), we set the relevant asymmetry parameter to 0.2, while keeping the other two asymmetry parameters fixed at 0. Fig. 4B shows snapshots of simulation outcomes for each case. A visual inspection suggests that these three cases give rise to a P compartments with similar cell sizes and shapes as in Fig. 3A.

Fig. 4C shows that P compartment sizes and cell numbers are not affected by these small amounts of asymmetry in the tissue. In each case, the *in silico* compartment area and cell number is as close to the observed values [14] as the passive mechanical model. Although cellular properties are now patterned, compartment size control still emerges within the model. Fig. 4D shows the total number of cell deaths in the front and the back halves of the P compartment across 100 simulations in each asymmetry case. We find that each case can explain the observed spatial asymmetry in cell death locations.

### Robustness of compartment size and compartment cell number to cellular asymmetry

To assess to which extent P compartment sizes and cell numbers are robust to spatial asymmetry in cell mechanical properties, we next vary each of the three asymmetry parameters in turn while keeping the others fixed at 0. Fig. 5 shows that increases in asymmetry lead to decreases in P compartment sizes and cell numbers (top and middle row) and the degree of asymmetry in cell death across the front and back halves of the compartment (bottom row).

In the model, P compartment sizes and cell numbers are most sensitive to asymmetry in cell target areas; for example, a value of *λ_A_* > 0.9 can result in loss of the entire P compartment. In contrast, P compartment sizes and cell numbers remain within experimentally measured regimes for values of *λ_p_* or *λ_l_* from 0 up to 0.4.

### Differential growth and mechanical regulation generate distinct distributions of cell shapes in *wt.*

To identify experimentally observable signatures to differentiate between modes of regulating compartment homeostasis, we examined the distributions of four measures of cellular morphology for the scenarios described in Fig. 4. We extract the distributions of cell areas, cell perimeters, lengths of edges between cells, and cell elongations within the P compartment at the end of each simulation. We observe distributions of these four measurements in the posterior compartment as a whole, and in the front and the back half of the compartment separately. The results of this investigation are summarized in Fig. 6.

The top two rows of Fig. 6 show that the distributions of cell areas and cell perimeters (row 1 and 2) for the area regulation scenario are distinct from the corresponding distributions for the line tension and the perimeter regulation scenario. In particular, the distribution of all areas is bimodal for the area regulation scenario, whereas it is not bimodal for the line tension and perimeter regulation scenarios. A similar distinction can be made for the perimeter distributions, which is bimodal for the ‘area regulation’ scenario and not bimodal for the ‘line tension regulation’ and ‘perimeter regulation’ scenarios. The bimodal distributions are marked by nearly non-overlapping distributions of cell areas and cell perimeters in the front and the back halves of the compartment for the area regulation scenario, whereas these distributions are overlapping in the line tension and perimeter regulation scenarios. Upon decomposing cell area distributions into contributions from the front and back halves of the P compartment, we see that the mean cell area is different between these two halves in the area and perimeter regulation scenarios, and the same holds for the cell perimeter distributions. Cell elongations and edge lengths have similar shapes and mean values for all three asymmetry scenarios (rows 3 and 4 of Fig. 6).

The results in Fig. 6 suggest that it is possible to distinguish between the ‘area regulation’ scenario (differential growth across the compartment) from the ‘line tension regulation’ scenario (regulation of apical mechanical properties) by measuring the distributions of cell areas or perimeters in the front and the back halves of the posterior compartment separately. The distribution of cell areas or perimeters across the P compartment may further allow one to distinguish the ‘area regulation’ scenario from the ‘perimeter regulation’ scenario, since this distribution is bimodal in the former scenario, but not clearly bi- or unimodal in the latter. However, multiple sources of noise in an experimental setup may make this distinction between the ‘area regulation’ and ‘perimeter regulation’ scenarios less clear. Measuring edge lengths or cell elongations will not reveal differences between the scenarios.

### Characteristics of cell area distributions for the *en>dap* and *en>CycE* perturbations are preserved across asymmetry scenarios

While cell area distributions in *wt* simulations may allow the different asymmetry scenarios considered to be distinguished from one another, these distributions in the *en>dap* and *en>CycE* cases provide model predictions that are preserved across all scenarios. In each case, we find that the cell area distribution is multimodal. In particular, the *en>dap* cell area distribution is trimodal in the ‘area regulation’ scenario, whereas it is bimodal in the other cases considered.

This multi-modality in areas arises from overlapping cell generations. Since we assume that cell target areas decrease upon division (*R*_*A*_ < 1), each successive generation of cells will have a smaller target area. In simulations of the *en> CycE* perturbation, some cells divide twice while others only divide once, resulting in a bimodal cell area distribution. Similarly, for the *en>dap* perturbation, some cells divide once while others don’t divide at all; however, we also observe area differences between cells in the front and the back half of the P compartment (Fig. 6). These effects combine to yield a trimodal cell area distribution.

In summary, the area distributions of the genetic perturbations *en>dap* and *en>CycE* may be used as a measure to validate the model assumptions, and provide a further tool to distinguish the ‘area regulation’ scenario from the ‘perimeter regulation’ and ‘line tension regulation’ scenarios.

### Simulated laser ablation experiments allow discrimination between asymmetry scenarios

As a further analysis of the model, we performed a laser ablation analysis on the final configuration of our *wt, en>dap* and *en>CycE* simulations. In 100 simulations for each perturbation, we ‘cut’ a randomly selected cell-cell interface (edge) in the P compartment. This was implemented by setting the line tension parameter 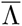 for this edge, as well as the perimeter tension parameter 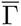 for the cells adjacent to the edge, to zero. We then ran each simulation for 200 further time units and recorded the average initial vertex recoil velocity and total vertex recoil distance. Results for each of the three asymmetry scenarios are shown in Fig. 8. We find that under the ‘perimeter regulation’ scenario, the average initial vertex recoil velocity and total vertex recoil distance are both smaller in each perturbation than in *wt.* In contrast, under the two other asymmetry scenarios there is no significant difference in these statistics across wt, *en>dap* and *en>CycE* simulations. These results offer a further experimentally testable prediction that, in conjunction with the cell area distribution results summarised in Fig. 6, allows for discrimination between the three asymmetry scenarios considered.

## Discussion

We have employed a vertex model of a *Drosophila* embryonic segment to test hypotheses about the emergence of size control. A comparison of the *in silico* segment with extant literature values indicated that passive mechanical forces suffice to explain the observed size control. However, the observed spatial heterogeneity in cell death frequencies requires some form of patterning of mechanical properties across the tissue. Several conceptually distinct modifications of the model can explain size control while also recapitulating the spatially varying rates of cell death: first, individual cells could regulate their sizes through differential growth; and second, cells could regulate their apical mechanical properties through differential expression of tension regulating protein activities. It is possible to distinguish these two scenarios within the model by the spatial distribution of P compartment cell areas and perimeters, as well as by the speed of vertex recoil after laser ablations. These results hint at two possible mechanistic functions of trophic signaling pathways, such as EGFR or Wg [14,36,37]: they could either cause growth of individual cells, or else modulate cell shape through regulation of contractile cytoskeletal activity, either of which would explain the experimentally observed shrinkage or growth when the pathways are perturbed [14].

### Connecting robustness of proportional size control to cell mechanics

Understanding the mechanism of tissue size control is particularly challenging due to the interconnected and complex nature of cell signaling and the high degree of feedback between cell- and tissue-level processes. Computational models therefore offer an important tool for investigating and testing hypothesised mechanisms and to abstract the principles underlying developmental robustness [38–40].

The development of multicellular organisms requires control of total cell numbers and relative proportions of cell types with tissues. Size control can be divided into two steps: initial specification and maintenance [12]. Much work has focused on the regulation of the position of cellular fates during early embryonic development. Traditionally, tissue size specification has been associated with signaling gradients [41–43]. However, the mechanisms that ensure the maintenance of tissue size and of boundaries between tissues is less well understood. In particular, the physics of size homeostasis for patterned epithelia are not well understood, yet they are a recurring theme in development [44, 45] and it is increasingly recognised that mechanical feedback plays a role in controlled tissue behaviour [8, 46].

A gradient growth model has previously been proposed for the regulation of P compartment size in the *Drosophila* embryonic epidermis [14]. This conceptual model requires the correct maintenance of a morphogen gradient in the face of multiple genetic perturbations. The present study demonstrates that an alternative, passive mechanical model can partially explain robustness of P compartment sizes and cell numbers in the *Drosophila* embryonic epidermis, eliminating the need for a tightly controlled intermediary morphogen gradient. More detailed cell-level analysis and modelling is required in the future to fully understand how morphogen signals are established, maintained, and interpreted [47, 48], especially in the face of genetic or environmental perturbations.

Advancing our knowledge of how embryos achieve robustness to defects or damage to the initial patterning of tissue domains is important for understanding the underlying causes of birth defects, as well as diseases with an underlying basis of misregulated growth, such as cancers.

### Providing predictions to guide future experimental inquiries into pattern repair

Although several studies have investigated the robustness of sizes of patterned epidermal segments of *Drosophila*, quantification has been somewhat sporadic and diffuse. This will in general require a thorough systems-level characterization of later stages of *Drosophila* morphogenesis for multiple experimental perturbations. The present study provides a basis for guiding future experiments that seek to identify possible modes of size control in late stages of epithelial development in *Drosophila.*

How could model predictions be validated against such experiments? Several previous vertex models of developing epithelia have been validated against key summary statistics. Such studies have focused primarily on the *Drosophila* wing imaginal disc, which undergoes up to 9 rounds of divisions to arrive at a distinct distribution of cell polygon numbers [22, 24]. In these studies, it is safe to assume that the initial distribution of cell polygon numbers will not affect simulation outcomes, due to the high levels of proliferation. Here, we considered one or two rounds of divisions; over such a short developmental timespan we expect the initial sheet topology to influence final polygon distributions. Hence, for a quantitative comparison of this summary statistic between model and data, experimentally informed cell shapes in late stage 10 segments may be required. Such summary statistics remain lacking for the *Drosophila* embryonic epidermis during its development, and poses an experimental challenge due to the small system size (20-60 cells). Large sample sizes will be required to obtain accurate distributions of cell polygon numbers. Figs. 7 and 8 in this study suggest that distributions of cell areas, and characterization of vertex recoils following standard laser ablation experiments, for genetic perturbations of the P compartment maybe used to validate the underlying computational model. Thus, future iterations of the model may be further constrained through inference of mechanical parameters from laser ablation [6] or less invasive experimental protocols [49].

**Figure 7.**
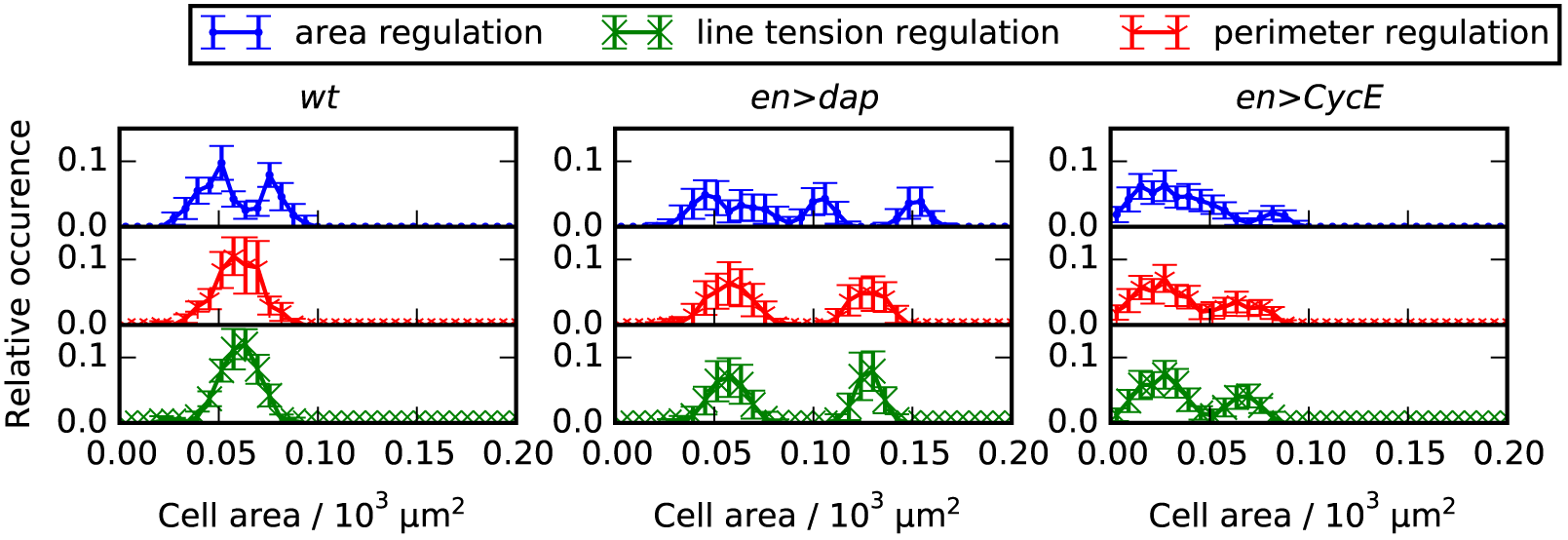
Cell area distributions in the *en>dap* and *en>CycE* perturbations are multimodal. Distributions of cell areas for each perturbation of cell division events (*wt, en>dap* and *en>CycE*) and each scenario of cellular asymmetry. Cell areas are recorded at the end of each simulation and error bars denote standard deviations across 100 simulations. Parameter values are given in Table 1 and in the main text.

**Figure 8.**
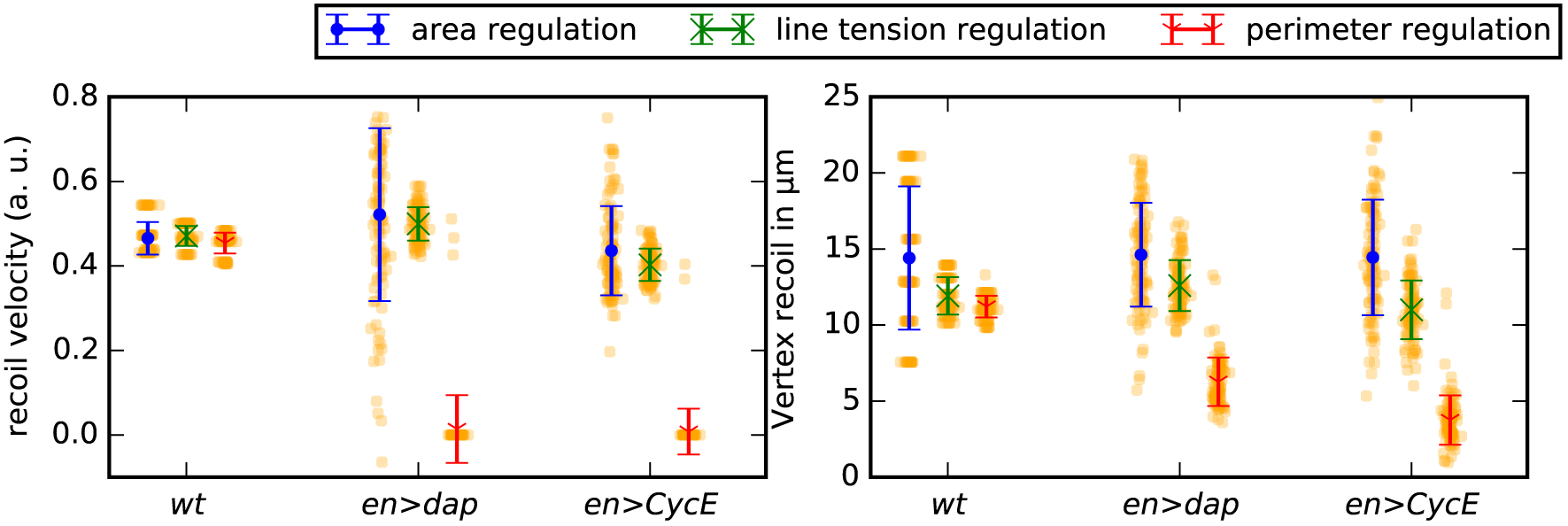
Simulated laser ablation experiments allow discrimination between asymmetry scenarios. Average initial vertex recoil velocities and total recoil distances across simulations of *wt* and perturbations. Error bars denote standard deviations across 100 simulations. Parameter values are given in Table 1 and in the main text.

### Current limitations

Embryogenesis is an extremely complex process. To make headway into the factors that influence robustness of tissue size maintenance, there needs to be conscious decoupling and abstraction through studies of simpler systems. This is also part of the rationale for studies in genetic model organisms from the worm and fly to mouse [40].

Due to the lack of kinematic data on cell shapes and compartment sizes during the latter stages of embryogenesis, we have not included an analysis for this initial study and have focused on more local mechanisms. In particular, we assumed that the overall tissue dimensions are constant during the considered time frame, since the epidermis forms at the outside of the embryo during stage five of *Drosophila* development and as a whole does not change dimensions for the remainder of development. However larger scale tissue morphogenetic movements, which are undoubtedly important for aspects of morphogenesis [50], may affect the exact size of a given subsection of the tissue. For example, dorsal closure occurs during the considered time frame, which leads to an extension of the tissue that we study [51]. The assumption that this extension should not affect the relative proportions in A and P compartment size requires future experimental validation. In addition, our finding that elevated tension along compartment boundaries does not affect compartment sizes may be contrasted with theoretical and experimental studies showing how differential line tension, either at compartment boundaries or across tissues, may drive convergent extension [52, 53]. A key conceptual difference between the present work and these studies is the assumption of a fixed, or free, boundary to the tissue.

In vertex models with a free boundary, contractile forces along cell perimeters may lead to deviations of cell areas from their respective target areas. The analysis of simulations with changed initial target areas presented in Fig. S4 reveals that such deviations between cell target areas and their absolute areas may lead to increases in predicted apoptotic rates. Further investigation is required to understand the boundary conditions that best represent the effect of adjacent tissues in different epithelia, and the effects that forces along tissue boundaries can have on different summary statistics. It may be possible to gain insights to this question by quantifying tissue-level kinematics of germ-band retraction for the *wt* and developmental perturbations.

Our model relies on the quasi-steady state assumption that the tissue is at mechanical equilibrium at each time point. We justified this assumption on the basis that individual cell cycle times of the 16th division cycle in *Drosophila* development are around an hour [9]. However, if cell divisions occur highly synchronously, then, this assumption might not hold. In *en>CycE* embryonic segments, the numbers of cell divisions events in the model were inferred from data where apoptosis was blocked by expressing the protein p35 in the P compartments [14]. It has previously been reported that epithelial sheets can extrude cells that are not undergoing apoptosis [29]; if this occurs to a great extent in the *Drosophila* embryonic epidermis, then our inferred numbers of mitotic events would require adjustment. In this case, an *in vivo* cell tracking study would be necessary to measure the levels of cell division and extrusion events. Such data would also help to shed light, for example, on the possible impact of mitotic cell rounding on local cell shapes and possible short-range correlations between mitosis and apoptosis events. Since apoptosis in the vertex model is a passive process, we cannot extend our model analysis to p35 mutants in which apoptosis is blocked. How to adapt vertex models in such a way as to prevent the occurrence of T2 transitions while remaining tissue integrity remains an open question.

Due to a current lack of data in the literature, our model does not include a description of upstream patterning of cell types. Instead, we infer the necessity of patterning of cell mechanics through simulations. This study is timely as it provides some guidance into important parameters and considerations that should be taken into account in future quantitative analyses of late stages of epidermal development including germband retraction and head involution. From the results presented here, further questions arise. If a passive mechanical model is sufficient to explain compartmental size control, then what is the functional role of Spitz-mediated EGFR regulation? It is known that EGFR signaling is required for dorsal closure during *Drosophila* development [54]. Hence, it is possible that the influence of EGFR signaling on larval compartment sizes reflects the role of EGFR signaling in convergent extension during dorsal closure. If the asymmetry in our model reflects patterning of mechanical properties through trophic signaling, then a more detailed experimental analysis of the spatio-temporal dynamics of cellular signaling will allow more detailed modelling of how these properties may be patterned.

### Summary and larger implications

Our study serves as an example of using computational models as an abstraction of the maintenance of tissue sizes with implications for a broad range of studies. Significant advances in stem cell engineering have resulted from understanding how to unlock the potential for multicellular aggregates to self-organize. Recent examples include the morphogenesis of optic eye cups in organ culture conditions [55] and the engineering of beating mini-hearts [56]. We posit that great success in developing tissue repair strategies will come through the reverse engineering of pattern repair mechanisms in situations where pattern repair is perturbed. Such reverse engineering will require guiding experimental efforts through modelling studies that identify the information needed to distinguish between mechanisms.

## Supporting Information

### Software S1

#### Zipped folder containing implementation of the computational model and analysis described in this study

A thoroughly documented example of how to run the code is provided in the file README.txt.

### Text S1

#### Immunohistochemistry

A *Drosophila* line expressing GAL4 under the engrailed (en) promotor and CD8::GFP under the UAS promotor was used. Immunohistochemistry on embryos was performed as described previously [57] with rabbit anti-dpERK (1:100, Cell Signaling), rat anti-DCAD2 (1:100, DSHB) and DAPI (5 μg/ml, Invitrogen DU1306), with goat anti-rat IgG 561 (1:500, Invitrogen), and goat anti-rabbit IgG 647 (1:500, Invitrogen).

#### Confocal microscopy

Confocal *z*-stacks were collected using an Andor spinning disc confocal microscope with a piezo stage at 1.0 μm intervals. For each embryo, six by six grids with thirty-three percent overlap were collected for each of the four channels. Image collection was performed with MetaMorph^®^ version 7.0.11. A costum vignetting correction algorithm was applied as a part of the stitching process.

#### Choice of simulation boundary condition does not significantly influence simulation results

In this section we analyse the influence of the choice of boundary condition on the compartment sizes and compartment cell numbers in the simulations. We compare the results of simulations using the doubly periodic boundary conditions described in the main text to those where fixed boundary conditions are imposed on the tissue as follows. Vertices on the boundary of the tissue experience the same forces as non-boundary vertices. At each time step, after applying the forces to all vertices and updating their positions, we map the boundary vertices perpendicularly back onto the boundary. If a cell on the boundary undergoes a T2 transition, then the newly created vertex is also mapped onto the boundary. The dimensions of this fixed boundary are chosen such that the total area of the tissue equals the sum of the target areas of all cells of the tissue. The tissue occupies a total of 64 rescaled area units, or 7.744×10^3^ μm^2^. The length of each fixed boundary is 8 *L*, i.e. 88 μm.

As Figure S1 shows, we find that imposing this alternative boundary condition gives similar predicted P compartment areas and cell numbers to those obtained using doubly periodic boundary conditions. This is true across *wt, en>dap* and *en>CycE* simulations.

#### Irregular initial conditions do not significantly influence simulation results

We next analyse the influence of the choice of initial tissue geometry on the final P compartment sizes and cell numbers in our model. In particular, we compare the results of simulations initiated with randomly generated cell shapes to those of simulations with regular hexagonal cell shapes. The random initial conditions are created as follows. First, 64 seeds are randomly placed in a spatial domain occupying 64 rescaled area units (7.744×10^3^ μm^2^). Second, the random seeds are mirrored along each tissue boundary to ensure periodicity, and the Voronoi tessellation of all seeds is computed. Third, we relax the Voronoi tessellation using five steps of Lloyd’s relaxation algorithm [58]. Cells are then assigned to the P compartment if their centroid falls within a central vertical band of width 2.9 rescaled length units (32 μm).

Figure S3 shows that these irregular initial conditions result in similar mean P compartment sizes and cell numbers to those obtained using our regular hexagonal initial conditions, while increasing the standard deviation in each quantity. Thus, irregularity in the initial geometric configuration does not significantly affect simulation outcomes.

#### Changes in initial target areas do not strongly affect simulation results

For both the boundary conditions considered in this study, the initial area of each cell *A^s^* takes its target value As 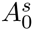. To test whether our results depend strongly on this assumption, we ran control simulations in which the initial area of each cell was half its target value at the beginning of the simulations. We found that resulting P compartment sizes and cell numbers were not strongly affected by this modification, and compartment size control was preserved. The results of this analysis are shown in Figure S4, and they differ from Fig. 3B in two ways. First, the tissue areas at the end of simulations vary slightly from the experimental observations made by Parker; in particular, the P compartment for the *en>cycE* perturbation is larger than for the *en>dap* perturbation. This is to be expected, since in this set of simulations most cell areas are far from their target values, hence the area energy term dominates and mitosis-induced shrinkage does not occur. Second, *en>CycE* P compartment cell numbers are smaller than observed by Parker and in the results of Fig. 3B (52 vs 60 and 72 cells, respectively), illustrating that more cells die in the simulations where cell target areas have been doubled. In the simulations summarized in Figure S4, the overall tissue target area is a lot larger than the tissue area, and therefore more cells are removed by apoptosis (T2 swaps) to decrease the corresponding energy term. Although we consider the initial condition for the simulations with doubled initial target areas As 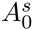 unrealistic, with cell areas being far from their target values, the model continues to exhibit P compartment size control.

**Table S1.**
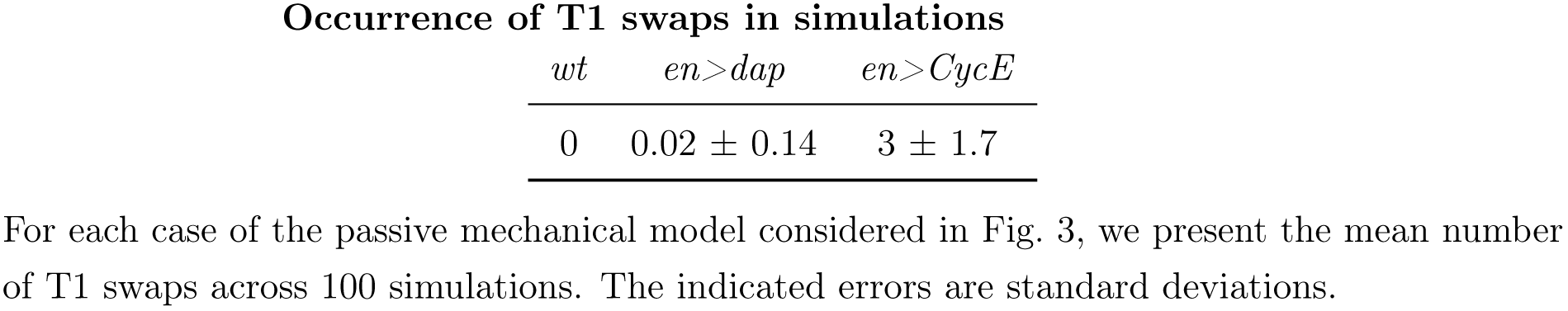
Occurrence of T1 swaps in simulations.

| **Occurrence of T1 swaps in simulations** |  |  |
| --- | --- | --- |
| *wt* | *en>dap* | *en>CycE* |
| 0 | $0.02 \pm 0.14$ | $3 \pm 1.7$ |

**Figure S1.**
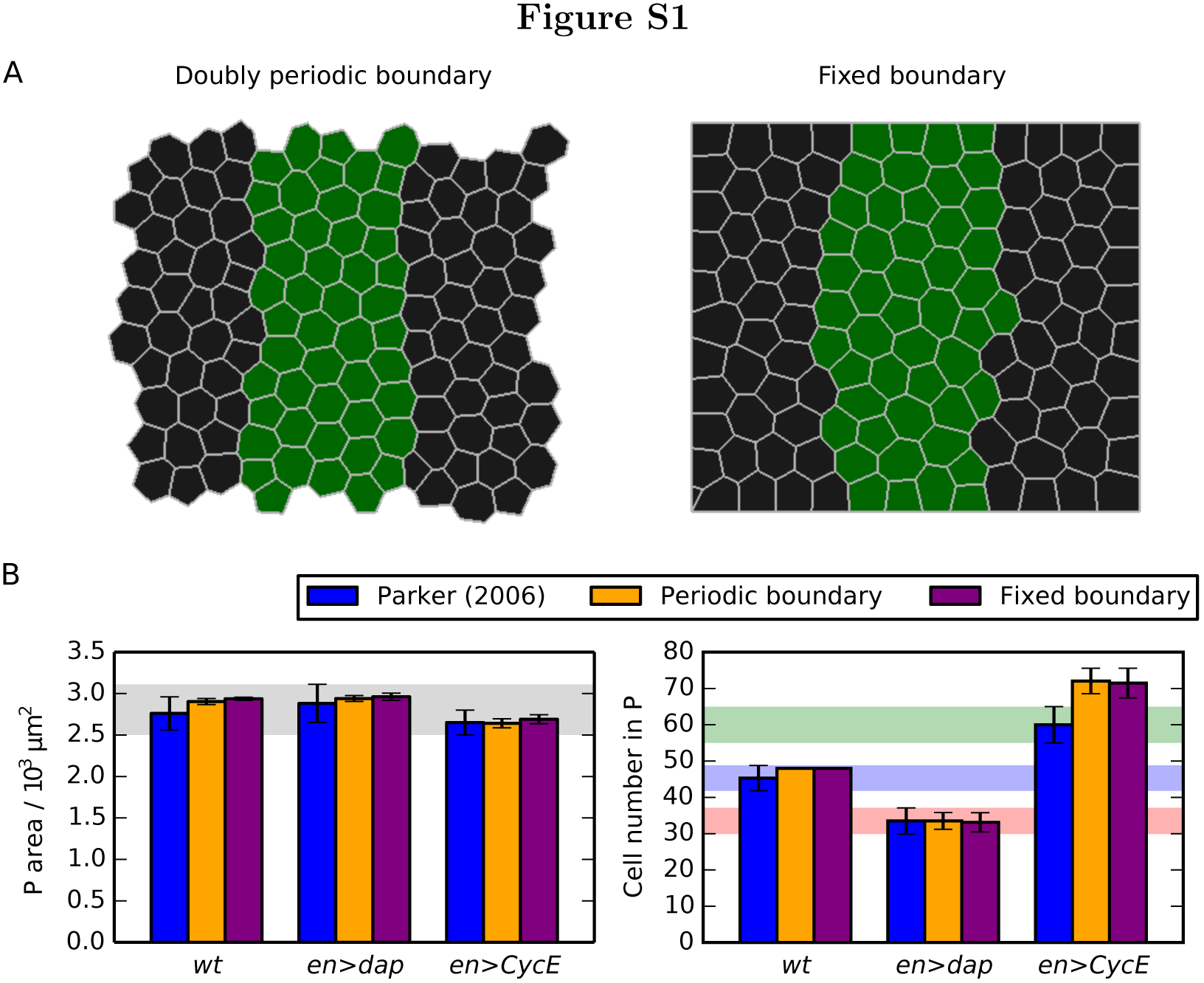
Choice of boundary condition does not affect P compartment sizes and cell numbers. (A) Snapshots of a *wt* simulation at the final time point, once all cell divisions have occurred, where doubly periodic (left) or fixed (right) boundary conditions are imposed. Parameter values are listed in Table 1. (B) Comparison of P compartment areas and cell numbers for *wt, en>dap* and *en>CycE* simulations where doubly periodic or fixed boundary conditions are imposed. Mean values from 100 simulations are shown and error bars are standard deviations. Shaded areas mark the ranges of experimentally observed values and are added for reference and comparison with Figs 3, 4 and S2.

**Figure S2.**
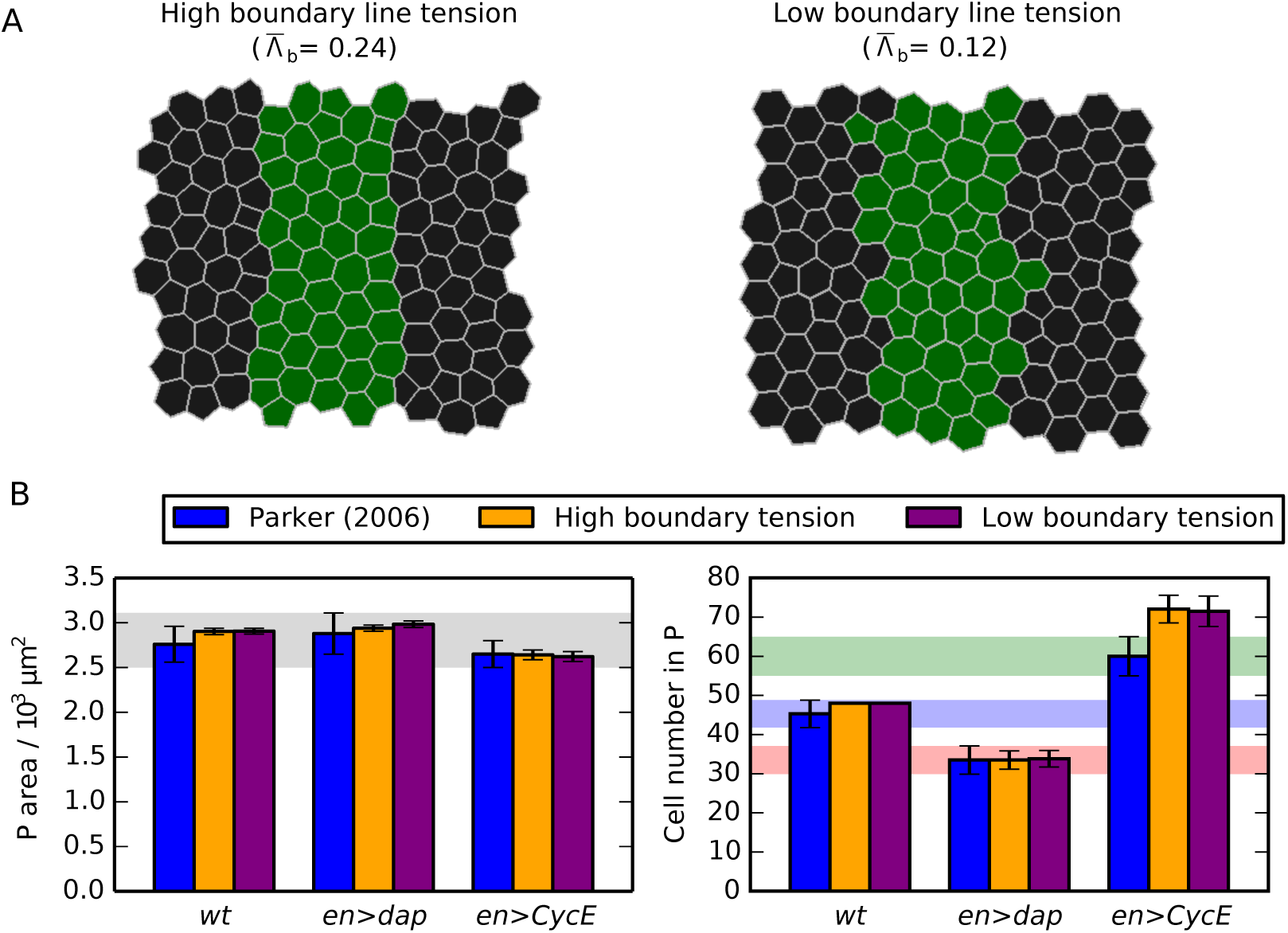
Compartment boundary line tension does not affect P compartment sizes and cell numbers. (A) Snapshots of a *wt* simulation at the final time point, once all cell divisions have occurred, where either a high (left) or low (right) line tension, 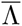, is imposed at the boundary between A and P compartments. Parameter values are listed in Table 1. Compartment boundary line tension promotes cell sorting and straightness of the boundary, but does not affect compartment sizes. (B) Comparison of P compartment areas and cell numbers for *wt, en>dap* and *en>CycE* simulations where a high (left) or low (right) compartment boundary line tension is imposed. Values for 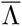 at the compartment boundary are those given in (A). Mean values from 100 simulations are shown and error bars are standard deviations. Shaded areas mark the ranges of experimentally observed values and are added for reference and comparison with Figs. 3, 4 and S1.

**Figure S3.**
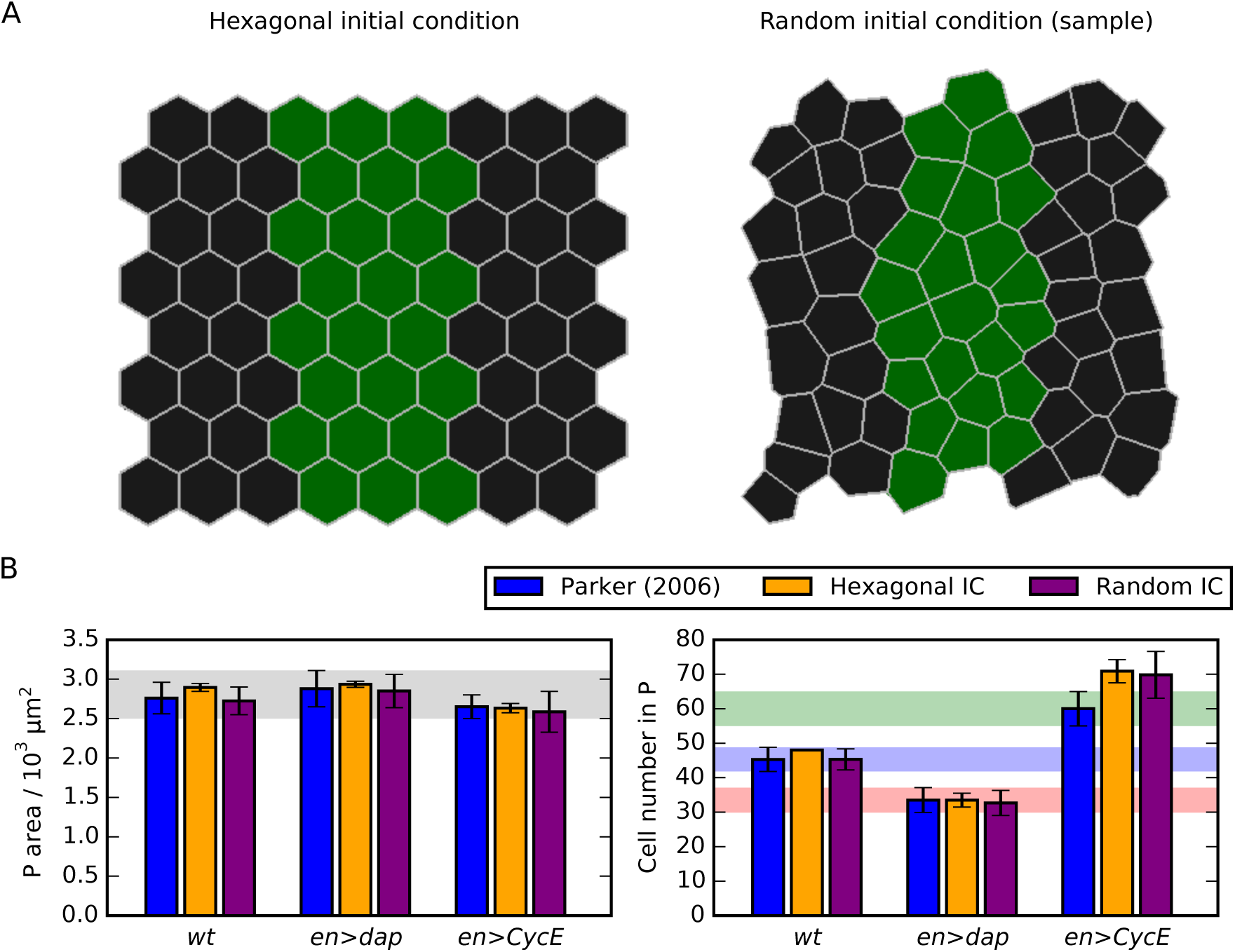
Initial cell shapes do not significantly affect P compartment sizes and cell numbers. (A) Snapshots of a hexagonal (left) initial condition, and a sample random (right) initial condition, as described in Text S1. The cells assigned to the posterior compartment occupy a similar area in both images. (B) Comparison of P compartment areas and cell numbers for *wt, en>dap* and *en>CycE* simulations where either a hexagonal or random initial condition (IC) was used. Mean values from 100 simulations are shown and error bars are standard deviations. Shaded areas mark the ranges of experimentally observed values and are added for reference and comparison with Figs. 3, 4 and S1. Parameter values are listed in Table 1.

**Figure S4.**
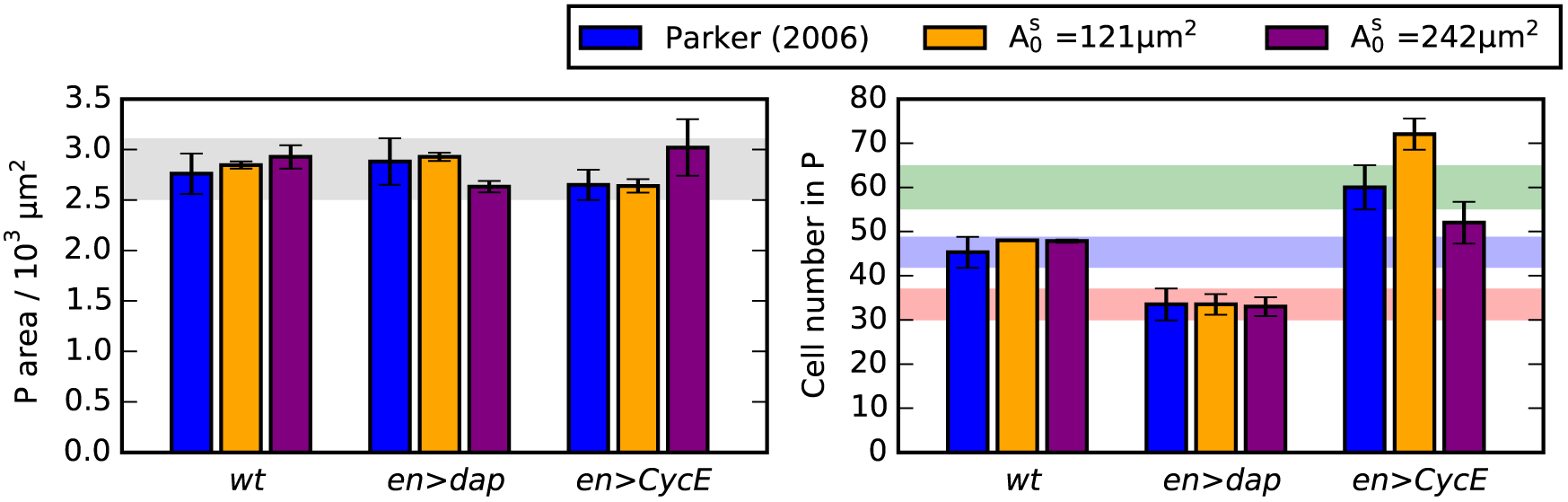
Influence of initial cell target area on P compartment size and cell numbers. (B) Comparison of P compartment areas and cell numbers for *wt*, *en>dap* and *en>CycE* simulations where either initial target areas 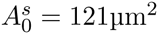 or 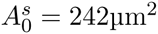 were used. Mean values from 100 simulations are shown and error bars are standard deviations. Shaded areas mark the ranges of experimentally observed values and are added for reference and comparison with Figs. 3, 4 and S1. Parameter values are listed in Table 1.

## Acknowledgments

JK acknowledges funding from the Engineering and Physical Sciences Research Council through a studentship. PB was funded partially by NIH grant UO1 HL116330. PB and JZ were funded in part by the National Science Foundation grant CBET 1403887. AF and JZ acknowledge funding from a Royal Society International Exchanges Scheme grant (IE130149). AGF acknowledges funding from the Engineering and Physical Sciences Research Council through grant EP/I017909/1 (www.2020science.net). PB and JZ acknowledge the Notre Dame Integrated Imaging Facility (NDIIF) to generate experimental figures. The authors would like to acknowledge the use of the University of Oxford Advanced Research Computing (ARC) facility in carrying out this work.

## References

1. Snow MHL, Tam PPL. Is compensatory growth a complicating factor in mouse teratology? Nature. 1979;279(5713):555–557.

2. Namba R, Pazdera TM, Cerrone RL, Minden JS. *Drosophila* embryonic pattern repair: how embryos respond to bicoid dosage alteration. Development. 1997;124(7):1393–1403.

3. Vernon JA, Butsch J. Effect of tetraploidy on learning and retention in the salamander. Science. 1957;125(3256):1033–1034.

4. Guillot C, Lecuit T. Mechanics of epithelial tissue homeostasis and morphogenesis. Science. 2013;340(6137):1185–1189.

5. Heisenberg CP, Bellaïche Y. Forces in tissue morphogenesis and patterning. Cell. 2013;153(5):948–962.

6. Landsberg KP, Farhadifar R, Ranft J, Umetsu D, Widmann TJ, Bittig T, et al. Increased cell bond tension governs cell sorting at the *Drosophila* anteroposterior compartment boundary. Curr Biol. 2009;19(22):1950–1955.

7. Monier B, Gettings M, Gay G, Mangeat T, Schott S, Guarner A, et al. Apico-basal forces exerted by apoptotic cells drive epithelium folding. Nature. 2015;518(7538):245–248.

8. Shraiman BI. Mechanical feedback as a possible regulator of tissue growth. Proc Natl Acad Sci USA. 2005;102(9):3318–3323.

9. Ashburner M. Drosophila: a laboratory handbook. 2nd ed. Cold Spring Harbor Press; 2011.

10. Kornberg T, Sidén I, O’Farrell P, Simon M. The engrailed locus of *Drosophila: In situ* localization of transcripts reveals compartment-specific expression. Cell. 1985;40(1):45–53.

11. Hughes SC, Krause HM. Establishment and maintenance of parasegmental compartments. Development. 2001;128(7):1109–1118.

12. Li QJ, Pazdera TM, Minden JS. *Drosophila* embryonic pattern repair: how embryos respond to cyclin E-induced ectopic division. Development. 1999;126(10):2299–2307.

13. Pazdera TM, Janardhan P, Minden JS. Patterned epidermal cell death in wild-type and segment polarity mutant *Drosophila* embryos. Development. 1998;125(17):3427–3436.

14. Parker J. Control of compartment size by an EGF ligand from neighboring cells. Curr Biol. 2006;16(20):2058–2065.

15. Perrimon N, Ni JQ, Perkins L. *In vivo* RNAi: today and tomorrow. CSH Perspect Biol. 2010;2(8):a003640.

16. Duffy JB. GAL4 system in *Drosophila*: a fly geneticist’s Swiss army knife. Genesis. 2002;34(1–2):1–15.

17. Elliott DA, Brand AH. The GAL4 system. In: Drosophila. Springer; 2008. p. 79–95.

18. Gavrieli Y, Sherman Y, Ben-Sasson SA. Identification of programmed cell death *in situ* via specific labeling of nuclear DNA fragmentation. J Cell Biol. 1992;119(3):493–501.

19. Foe VE. Mitotic domains reveal early commitment of cells in *Drosophila* embryos. Development. 1989;107:1–22.

20. Song Z, McCall K, Steller H. DCP-1, a *Drosophila* cell death protease essential for development. Science. 1997;275(5299):536–540.

21. Nagai T, Kawasaki K, Nakamura K. Vertex dynamics of two-dimensional cellular patterns. J Phys Soc Jpn. 1988;57(7):2221–2224.

22. Farhadifar R, Röper JC, Aigouy B, Eaton S, Jülicher F. The influence of cell mechanics, cell-cell interactions, and proliferation on epithelial packing. Curr Biol. 2007;17(24):2095–2104.

23. Nagai T, Honda H. A dynamic cell model for the formation of epithelial tissues. Philos Mag B. 2001;81(7):699–719.

24. Aegerter-Wilmsen T, Smith AC, Christen AJ, Aegerter CM, Hafen E, Basler K. Exploring the effects of mechanical feedback on epithelial topology. Development. 2010;137(3):499–506.

25. Mao Y, Tournier AL, Hoppe A, Kester L, Thompson BJ, Tapon N. Differential proliferation rates generate patterns of mechanical tension that orient tissue growth. EMBO J. 2013;32(21):2790–2803.

26. Fletcher AG, Osterfield M, Baker RE, Shvartsman SY. Vertex models of epithelial morphogenesis. Biophys J. 2014;106(11):2291–2304.

27. Honda H, Nagai T. Cell models lead to understanding of multi-cellular morphogenesis consisting of successive self-construction of cells. J Biochem. 2015;157(3):129–136.

28. Schöck F, Perrimon N. Cellular processes associated with germ band retraction in drosophila. Dev Biol. 2002;248(1):29–39.

29. Marinari E, Mehonic A, Curran S, Gale J, Duke T, Baum B. Live-cell delamination counterbalances epithelial growth to limit tissue overcrowding. Nature. 2012;484(7395):542–545.

30. Canela-Xandri O, Sagués F, Casademunt J, Buceta J. Dynamics and mechanical stability of the developing dorsoventral organizer of the wing imaginal disc. PLoS Comput Biol. 2011;7(9):e1002153.

31. Fletcher AG, Osborne JM, Maini PK, Gavaghan DJ. Implementing vertex dynamics models of cell populations in biology within a consistent computational framework. Prog Biophys Mol Biol. 2013;113(2):299–326.

32. Hofmeister W. Zusätze und Berichtigungen zu den 1851 veröffentlichen Untersuchungengen der Entwicklung höherer Kryptogamen. Jahrb wiss Bot. 1863;3:259–293.

33. Gibson WT, Veldhuis JH, Rubinstein B, Cartwright HN, Perrimon N, Brodland GW, et al. Control of the mitotic cleavage plane by local epithelial topology. Cell. 2011;144(3):427–438.

34. Monier B, Pelissier-Monier A, Brand AH, Sanson B. An actomyosin-based barrier inhibits cell mixing at compartmental boundaries in *Drosophila* embryos. Nat Cell Biol. 2010;12(1):60–65.

35. Mirams GR, Arthurs CJ, Bernabeu MO, Bordas R, Cooper J, Corrias A, et al. Chaste: an open source C++ library for computational physiology and biology. PLoS Comput Biol. 2013;9(3):e1002970.

36. O’Keefe L, Dougan ST, Gabay L, Raz E, Shilo BZ, DiNardo S. Spitz and Wingless, emanating from distinct borders, cooperate to establish cell fate across the Engrailed domain in the Drosophila epidermis. Development. 1997;124(23):4837–4845.

37. Szuts D, Freeman M, Bienz M. Antagonism between EGFR and Wingless signalling in the larval cuticle of Drosophila. Development. 1997;124(16):3209–3219.

38. Iber D, Tanaka S, Fried P, Germann P, Menshykau D. Simulating tissue morphogenesis and signaling. In: Tissue Morphogenesis. Springer; 2015. p. 323–338.

39. Schwarz US, Dunlop CM. Developmental biology: a growing role for computer simulations. Curr Biol. 2012;22(11):R441–R443.

40. Xiong F, Megason SG. Abstracting the principles of development using imaging and modeling. Integr Biol. 2015;7:633–642.

41. Wolpert L. Positional information and the spatial pattern of cellular differentiation. J Theor Biol. 1969;25(1):1–47.

42. Crick F. Diffusion in embryogenesis. Nature. 1970;225(5231):420–422.

43. Rogers KW, Schier AF. Morphogen gradients: From generation to interpretation. Annu Rev Cell Dev Biol. 2011;27(1):377–407.

44. Vakulenko S, Reinitz J, Radulescu O. Size regulation in the segmentation of drosophila: interacting interfaces between localized domains of gene expression ensure robust spatial patterning. Phys Rev Lett. 2009;103(16):168102.

45. von Dassow M, Davidson LA. Physics and the canalization of morphogenesis: a grand challenge in organismal biology. Phys Biol. 2011;8(4):045002.

46. Buchmann A, Alber M, Zartman JJ. Sizing it up: the mechanical feedback hypothesis of organ growth regulation. In: Semin. Cell Dev. Biol.. vol. 35; 2014. p. 73–81.

47. Hufnagel L, Teleman AA, Rouault H, Cohen SM, Shraiman BI. On the mechanism of wing size determination in fly development. Proc Natl Acad Sci USA. 2007;104(10):3835–3840.

48. Wartlick O, Mumcu P, Jülicher F, González-Gaitán M. Understanding morphogenetic growth control - lessons from flies. Nat Rev Mol Cell Biol. 2011;12(9):594–604.

49. Ishihara S, Sugimura K. Bayesian inference of force dynamics during morphogenesis. J Theor Biol. 2012;313(0):201–211.

50. Butler LC, Blanchard GB, Kabla AJ, Lawrence NJ, Welchman DP, Mahadevan L, et al. Cell shape changes indicate a role for extrinsic tensile forces in drosophila germ-band extension. Nat Cell Biol. 2009;11(7):859–864.

51. Gorfinkiel N, Blanchard GB, Adams RJ, Martinez Arias A. Mechanical control of global cell behaviour during dorsal closure in drosophila. Development. 2009;136(11):1889–1898.

52. Rauzi M, Verant P, Lecuit T, Lenne PF. Nature and anisotropy of cortical forces orienting drosophila tissue morphogenesis. Nat Cell Biol. 2008;10(12):1401–1410.

53. Vroomans RM, Hogeweg P, ten Tusscher KHWJ. Segment-specific adhesion as a driver of convergent extension. PLoS Comput Biol. 2015;11(2):e1004092–e1004092.

54. Shen W, Chen X, Cormier O, Cheng DC, Reed B, Harden N. Modulation of morphogenesis by EGFR during dorsal closure in *Drosophila*. PLoS ONE. 2013;8(4):e60180.

55. Eiraku M, Takata N, Ishibashi H, Kawada M, Sakakura E, Okuda S, et al. Self-organizing optic-cup morphogenesis in three-dimensional culture. Nature. 2011;472(7341):51–56.

56. Ma Z, Wang J, Loskill P, Huebsch N, Koo S, Svedlund FL, et al. Self-organizing human cardiac microchambers mediated by geometric confinement. Nat Commun. 2015;6(7413).

57. Zartman JJ, Kanodia JS, Cheung LS, Shvartsman SY. Feedback control of the EGFR signaling gradient: superposition of domain-splitting events in *Drosophila* oogenesis. Development. 2009;136(17):2903–2911.

58. Lloyd S. Least squares quantization in PCM. IEEE Trans Inform Theory. 1982;28(2):129–137.

